# Localising the Seizure Onset Zone from Routine Interictal and Ictal Scalp EEG Via a Factorised Spatiotemporal Ensemble Deep Learning Framework

**DOI:** 10.64898/2026.10.16.759501

**Authors:** Songyuan Shi

**Affiliations:** Department of Computational Neuroscience, University College London, London, UK; Department of Biomedical Engineering, King’s College London, London, UK

**Keywords:** Seizure Onset Detection, Interictal Discharges, Transformer Neural Network, Non-invasive Diagnostics, Clinical Decision Support, Lobar Classification, Presurgical Screening

## Abstract

**Background:** Drug-resistant focal epilepsy requires precise localisation of the epileptogenic zone (EZ) for surgical resection. Current practice depends on invasive recordings or spontaneous seizure capture, a costly and risky bottleneck. Interictal scalp EEG can exhibit localising features like spikes, but scalp EEG suffers from volume conduction (the smearing of brain signals across multiple electrodes) and low signal-to-noise ratio (SNR), which obscure subtle spatial dynamics. Neurophysiologist review of long, often multi-day recordings is time-consuming and expertise-dependent, and while computational methods exist, they tend to rely heavily on human-designed features. Deep learning for EEG localisation could therefore help to improve clinical efficiency and sensitivity.

**Objectives:** To develop and validate a non-invasive deep learning approach for lobar-level localisation of the seizure onset zone (SOZ), the primary electrophysiological proxy for the epileptogenic zone (EZ), using interictal and ictal segments from routine scalp EEG. The model is trained on expert-annotated clinical labels and is intended to augment, not supplant, clinical judgement in EZ delineation.

**Methods:** We curated 262 sessions (6,000+ hours) from 100 patients with unilateral focal epilepsy, tested with an extra 28-patient internal and a 9-patient external dataset. The framework integrates: (1) an online augmentation pipeline (Gaussian noise, amplitude scaling, temporal cropping, and hemispheric channel swap) to mitigate overfitting. (2) a multi-scale temporal block for 20-200 ms discharge morphologies; (3) a factorised spatiotemporal Transformer decoupling temporal and spatial attention to counter volume conduction; and (4) a hemispheric mutual exclusivity penalty encoding unilateral priors. A five-model heterogeneous ensemble, fused via model soup, was evaluated on held-out, seizure-free, and cross-institutional cohorts.

**Results:** The ensemble achieved 88.46% session-level exact match on internal test data, outperforming EEGNet (50.00%). Performance generalised to the seizure-free holdout set (82.76%) and to cross-institutional data (84.62%). Rare occipital class (RO) recall rose from 0.500 to 1.000, with zero false positives on healthy controls (n=108). On unlabelled random segments, accuracy was 66.67% (vs. 14% chance). A full 24-hour recording was processed in 4.26 minutes, with inference under 8 seconds, highlighting localising windows for targeted expert clinical review

**Conclusion:** Clinically actionable SOZ spatial information is encoded in routine interictal scalp EEG. Our factorised ensemble provides a safe, interpretable, and efficient tool to expedite presurgical evaluation, potentially reducing reliance on prolonged monitoring and invasive implantation.

## 1: Introduction

### 1.1 Clinical Background, Presurgical Localisation Bottlenecks and EEG Electrodes Arrangement

Epilepsy is a chronic neurological condition defined by recurrent, unprovoked seizures caused by abnormal, excessive electrical activity in the brain. Epilepsy affects over 50 million individuals globally [1], and its clinical manifestations exhibit marked heterogeneity, spanning from brief lapses of awareness to generalised tonic-clonic convulsions that incur severe risks of secondary physical trauma. Although first-line anti-seizure medications (ASMs) are widely accessible, about 30% to 50% of patients with focal epilepsy do not respond to medications, failing to achieve long-term seizure freedom even with multiple drug combinations [2]. For this cohort with drug-resistant epilepsy (DRE), surgical resection or disconnection of the epileptogenic tissue offers the only viable therapeutic options to eliminate seizures and restore neurological function.

From a functional neurosurgical perspective, long-term postoperative seizure freedom relies on the precise and complete resection of the epileptogenic zone (EZ) - meaning the minimum brain tissue that must be removed to stop seizures [3]. Because the EZ cannot be directly visualised preoperatively, clinicians rely on observable biomarkers to delineate its boundaries. In current presurgical evaluation workflows, electroencephalogram (EEG), semiology and magnetic resonance imaging (MRI) are all used for seizure localisation; MRI can localise macroscopic morphological abnormalities, such as focal cortical dysplasia (FCD), hippocampal sclerosis, low-grade neuroepithelial tumours, and vascular malformations. However, up to 30% of focal DRE cases present as MRI-negative (non-lesional epilepsy) [4], showing no detectable structural lesions under conventional 3T imaging. Even in MRI-positive cases, the anatomical boundary of a structural lesion may mismatch with the true dynamic boundary of the epileptogenic network [5]. In these cases, static structural imaging fails to capture the spatiotemporal evolution of paroxysmal hypersynchronous electrical activity.

EEG offers a different functional perspective by providing real-time brain electrical signals with high temporal resolution. Modern VEEG combines both EEG recording and video recording, which allows clinicians to match brain wave abnormalities with clinical manifestations and enables the precise tracking of ictal activity from its spatial point of origin (the seizure onset zone (SOZ)), through its propagation across specific neural circuits, to its eventual termination. This analysis provides critical functional spatial guidance for surgical margin demarcation [6]. Nevertheless, classical non-invasive localisation methods are bottlenecked by their dependency on capturing true ictal (seizure) events [7]. Capturing spontaneous seizures typically requires prolonged, costly inpatient monitoring spanning several days to weeks, involves tapering anti-seizure medications, and exposes patients to heightened risks. Therefore, predicting the spatial location of the SOZ from interictal data can circumvent some hazards of ictal recording.

In this study, the International 10-20 System (Figure 1) was applied to ensure consistent electrode placement across all subjects, regardless of head size. For feature extraction, we converted the data into an 18-channel bipolar montage (the “Double Banana” configuration, Figure 1) [8] in this study.

**Figure 1:**
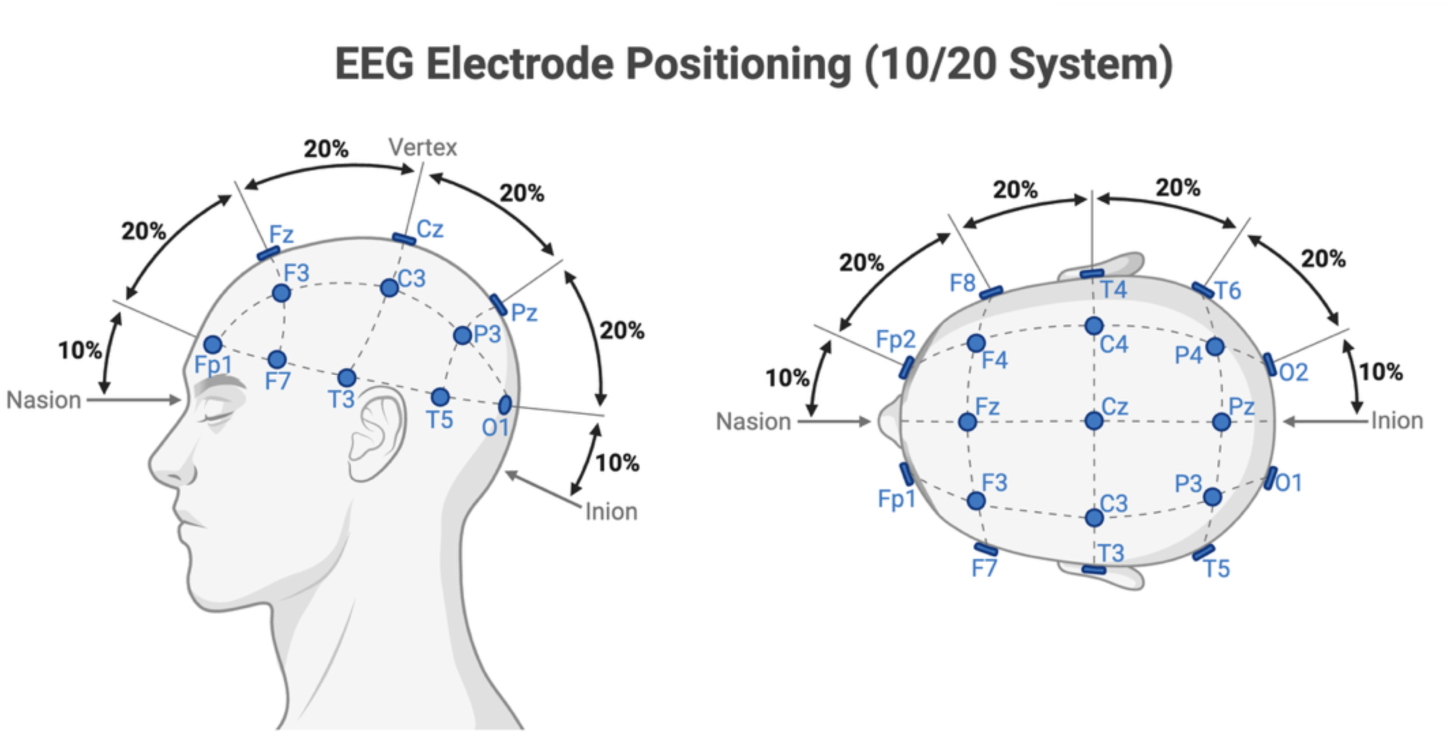
Standard 10–20 system electrode placement. Scalp EEG electrodes were positioned according to the International 10–20 System, ensuring reproducible spatial alignment across subjects regardless of variations in head circumference. The montage includes 19 standard electrodes: Fp1/Fp2 (frontal polar), F7/F3/Fz/F4/F8 (frontal), T3/C3/Cz/C4/T4 (central/temporal), T5/P3/Pz/P4/T6 (parietal), and O1/O2 (occipital). Nasion and inion serve as anatomical landmarks for midline alignment. This standardised positioning provides the spatial framework for subsequent bipolar longitudinal montage derivation and six-region anatomical classification.

**Figure 2:**
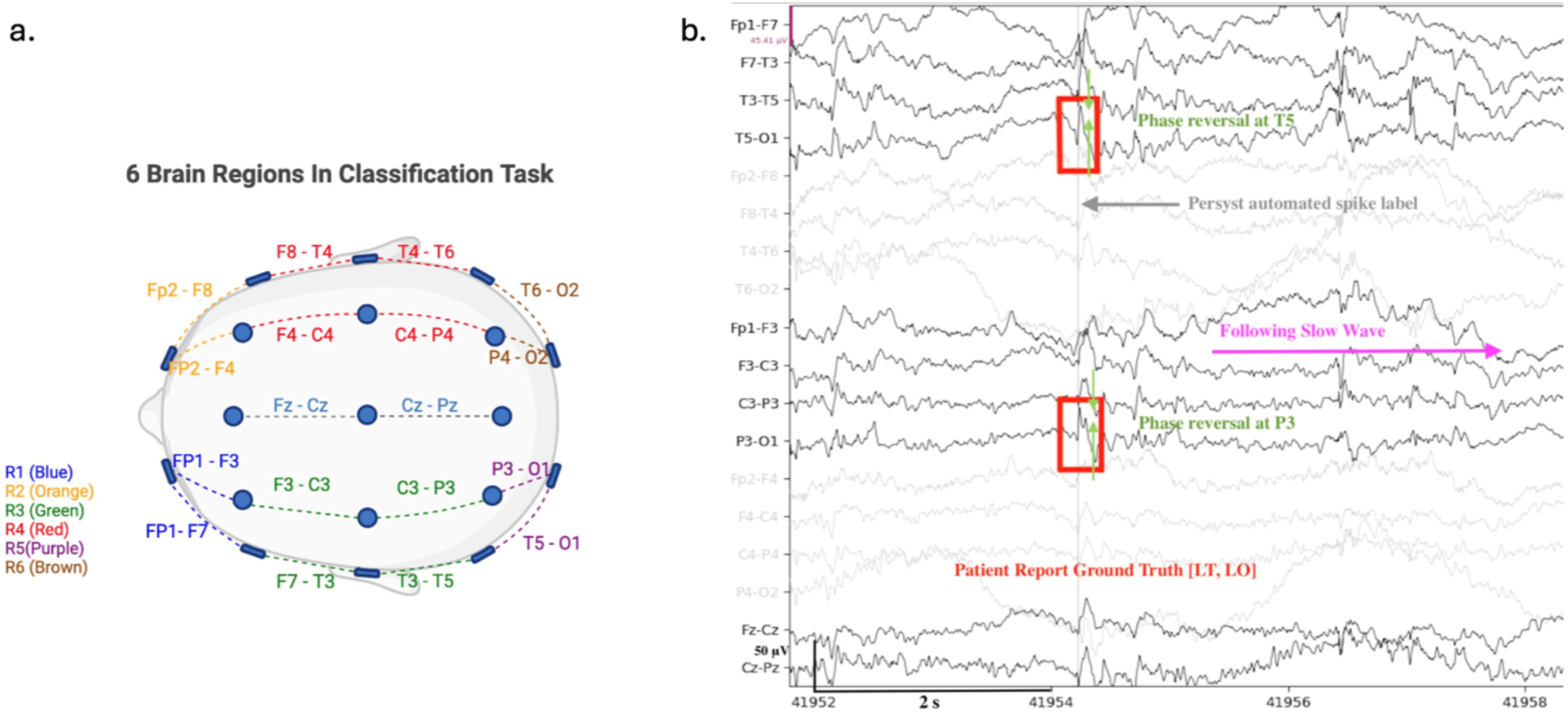
(a)Six-region anatomical mapping for the classification task. The 18 bipolar EEG channels are segmented into six brain regions corresponding to primary cortical lobes: Left Frontal, blue, Right Frontal, orange, Left Temporo-Parietal, green, Right Temporo-Parietal, red, Left Occipital, purple, and Right Occipital, brown. Temporal and parietal lobes are combined into single regions due to the spatial resolution limits of the bipolar longitudinal montage (inter-electrode distance 6–8 cm), which blurs the boundary between adjacent temporal (T3/T4) and parietal (P3/P4) electrode pairs. (b)A raw EEG annotated segment coming from the training set with Persyst automated spike labels; the ground truth is the left temporal and occipital region SOZ, This panel shows how spikes and phase reversals look clinically and the follow-up slow wave.

Then the raw scalp potentials were re-referenced using a bipolar longitudinal montage (e.g., by calculating the differential-voltage channel signal as Fp1 - F7) [8]. This differential approach filters out common-mode artefact noise and amplifies localised, high-frequency interictal transients. The downstream task is configured based on this arrangement as a lobe-level 6-class classification. The channels are segmented into six distinct anatomical regions mapping directly to primary cortical lobes: Left Frontal, Right Frontal, Left Temporal/Parietal, Right Temporal/Parietal, Left Occipital, and Right Occipital.

This mapping into six regions is not arbitrary but is a direct consequence of the resolution limits of the clinical imaging protocol. In standard scalp EEG, the difference in voltage between adjacent electrodes attenuates far-field activity and amplifies localised transients. However, spatial smoothing of the bipolar derivation with the 6–8 cm inter-electrode distance hampers accurate discrimination of neighbouring temporal and parietal electrode pairs, resulting in a blurred boundary between the neocortical regions. Also, pure parietal region seizures are very rare; virtually all patients with parietal seizures also have seizure onset in the temporal region; thus, separating the temporal and parietal lobes into different classes may cause the model to learn to adapt to noise from clinical scalp EEG, resulting in severe overfitting and poor generalisation across sessions. From a clinical workflow perspective, this lobe-level classification serves as a front-line triage tool: it identifies the dominant macro-region of epileptogenicity, thereby guiding clinicians toward the most probable targets for subsequent stereotactic intracranial electrode (sEEG) implantation [9], where finer sublobar discrimination can be achieved invasively.

### 1.2 Scientific Foundation of Interictal Spike Localisation

Since the ictal data are scarce, recent research has increasingly shifted toward interictal epileptiform discharges (IEDs) to circumvent the dependence on capturing such events. Pathophysiologically, interictal spikes are brief bursts of highly synchronised neuronal activity arising from the epileptic network and therefore offer promise for localising the epileptogenic tissue. Interictal discharges exhibit substantial morphological heterogeneity: brief spikes (20–70 ms), sharp waves (70–200 ms), and slow-wave complexes (4–12 Hz) are all epileptiform biomarkers with distinct pathophysiological generators [10].

Recent quantitative studies have demonstrated the spatial stability and clinical reliability of these interictal spikes. For instance, Jiang et al. [11] showed, through a spatial–temporal– spectral imaging (STSI) framework, that tracking these biomarker configurations yields a stable spatial projection of the epileptogenic tissue, achieving a mean localisation error of 19.59 mm relative to the surgically resected areas. Their data also confirmed that analysing approximately 30 to 40 high-quality interictal spikes is sufficient to yield stable, reliable localisation results, thereby significantly reducing the clinical reliance on capturing prolonged full ictal events. This is consistent with recent advancements in high-density scalp electroencephalography (EEG) source imaging; for example, the DeepSIF framework developed by Sun et al. [12] demonstrated that deep learning can improve source localisation accuracy relative to beamforming on high-density EEG.

Conventional EEG source imaging (ESI) techniques, such as sLORETA, MNE, and dSPM, have delivered high diagnostic utility under optimal settings, achieving a sensitivity of 84% and a specificity of 88% in identifying the irritative zone among surgical patients [13]. However, these physics-based methods construct individual head models from MRI scans and rely on high-density electrode arrays, precise 3D electrode digitisation, and subject-specific structural MRI for accurate volume conductor modelling [14]; these requirements are often unavailable or impractical in standard clinical routines. Their heavy reliance on individual anatomy and computational inverse solvers fundamentally limits their scalability to routine clinical recordings, where head models are rarely available, and electrode placement may vary across sessions.

Volume conduction refers to the passive spread of electrical potentials through the skull, scalp, and cerebrospinal fluid, and blurs the spatial specificity of scalp EEG, such that a single cortical source projects diffusely across multiple electrodes [23]. This physical constraint fundamentally limits the spatial resolution of scalp recordings regardless of electrode density.

In contrast, an end-to-end deep learning framework learns the inverse mapping directly from raw data, bypassing explicit forward-model assumptions while preserving single-spike temporal resolution. Therefore, our framework is not intended to replace high-density ESI for millimetre-level sublobar localisation, but rather a practical tool for standard clinical use. Specifically, this study aims to generate pilot evidence that a deep learning framework for SOZ localisation, operating without predetermined feature selection, can effectively localise the clinically defined seizure onset zone.

It is important to distinguish the present work from the substantial body of literature employing intracranial EEG (iEEG) for SOZ localisation. While iEEG offers unparalleled signal-to-noise ratio and direct cortical access, its invasive nature, requiring stereotactic implantation with attendant risks of haemorrhage and infection [15] renders it a secondary procedure reserved for patients who have already exhausted non-invasive evaluation. From a clinical workflow perspective, the rate-limiting step is not the analysis of iEEG data, but the initial decision of where to implant the electrodes, a decision currently guided by scalp EEG, neuroimaging, and semiology [9]. Therefore, enhancing the spatial informativeness of routine scalp EEG has the potential to impact a much larger patient population by reducing the number of patients requiring invasive monitoring and guiding more accurate implantation when iEEG is ultimately necessary.

### 1.3 Evolution and Limitations of Machine Learning in Seizure Onset Zone Localisation

Given the clinical value of interictal spikes, the computational neuro-medicine domain has seen an influx of automated frameworks leveraging machine learning (ML) and deep learning (DL) to localise the SOZ. The technological evolution of these algorithms can be characterised by two primary phases, each presenting distinct computational and clinical bottlenecks.

The early research heavily relied on traditional machine learning frameworks utilising hand-crafted feature extraction. Methodologies frequently extracted time- and frequency-domain metrics, such as information entropy, fractal dimensions, and power spectral density, from digitised EEG signals. Researchers subsequently deployed classical classifiers, including Support Vector Machines (SVM) and Random Forests [16] to perform automated epileptic seizure detection during ictal periods [17]. Although these classic models showed high Proficiency in detecting the onset of temporal ictal events within tightly controlled environments, their paradigms were fundamentally designed for transient wave detection rather than spatial source mapping. They also exhibit poor generalisability in real-world clinical settings [17] because they rely on rigid, handcrafted features that fail to accommodate the profound inter-patient and inter-session variability inherent in clinical EEG recordings.

There have been many deep learning approaches applied to EEG and epilepsy, but these have predominantly focused on temporal seizure detection with very little on spatial SOZ localisation, leaving this area largely underexplored. The conceptual framework of generic convolutional architectures limits their direct scalability to clinical EZ localisation tasks.

Generic pipelines, such as EEGNet, ShallowConvNet, and DeepConvNet, were originally developed for macro-level cognitive EEG decoding or transient seizure-pattern classification [18, 19]. This fundamental task mismatch creates a major architectural limitation when shifting from detection to localisation: these models utilise fixed, broad receptive fields optimised for extracting global temporal variations, making them inherently ill-suited to capturing the localised, micro-level morphologies of scalp spikes (20–200 ms) required for source mapping. As a result, while they may achieve satisfactory benchmarks in temporal gating or classification, they lack the necessary spatial inverse-mapping capacity to serve as standalone EZ localisation tools in routine clinical environments. Because ictal scalp recordings are systematically contaminated by severe myographic and motion artefacts induced by patient convulsions, relying solely on these detection-orientated networks for spatial inference often leads to elevated false-positive rates on cross-centre clinical data. Therefore, a key methodological gap remains between temporal pattern detection and precise spatial source localisation.

Subsequently, advanced deep learning architectures, such as Graph Convolutional Networks (GCNs), were introduced to model the brain as a dynamic network topology for non-invasive mapping. By integrating EEG source imaging (ESI) with geometric deep learning, these frameworks project sensor-level scalp recordings into the cortical source space, constructing graph representations where nodes represent reconstructed cortical regions rather than raw electrode channels [21]. Although these source-space architectures have achieved high area-under-the-curve metrics (AUC > 0.90) on small-scale datasets, their clinical deployment faces three key challenges:

**Challenge 1—Intracranial data bias:** Most high-performance models are trained on invasive EEG (iEEG), which offers high signal quality but limited spatial coverage. This does not reflect the reality of scalp EEG—the tool used in initial patient screening—and introduces a significant source of bias for data-driven models [13].
**Challenge 2—Task oversimplification:** Most existing paradigms treat epilepsy evaluation as a temporal event detection task (e.g., detection of spikes or seizures within isolated epochs) rather than explicitly modelling full-brain interictal dynamics or mapping sensor-level features to the SOZ location.
**Challenge 3—Small sample sizes:** Most published models are trained on less than 30 patients leading to patient-specific overfitting. This is compounded on scalp EEG, which suffers from volume conduction and spatial smearing [23]; networks trained without robust constraints rapidly degrade on cross-centre data.

Class imbalance presents another challenge in epileptic EEG datasets. Seizure periods are intrinsically rare relative to interictal periods, and certain epilepsy subtypes, such as occipital lobe epilepsy, are epidemiologically under-represented. This compound imbalance biases learning algorithms toward majority classes, leading to models that perform well on normal EEG but fail to detect subtle spatial or temporal variations. However, how to mitigate this dual imbalance, which exists both temporally (seizure vs. non-seizure) and spatially (across asymmetrical anatomical regions), has received little attention in multi-label SOZ localisation. Our framework tackles this issue by combining focal loss with region-specific positive weighting targeted for rare occipital classes.

Several recent studies have advanced SOZ localisation beyond earlier CNN- and GCN-based approaches. DeepSOZ [24] employs a Transformer encoder with global and channel-wise encodings combined with attention-weighted multi-instance pooling, validated on 120 patients from the Temple University Hospital corpus, yet relies on high-density scalp EEG. Ntolkeras et al. [25] applied machine learning to interictal EEG source connectivity in paediatric drug-resistant epilepsy. For invasive recordings, SEEGformer [26], a Transformer trained on 61 patients across two centres, was proposed for seizure detection and EZ localisation using stereo-EEG data. More recently, MBAT (Multi-Band Adaptive Transformer) [27] has been proposed for joint seizure detection and SOZ localisation from scalp EEG. However, these approaches predominantly rely on either invasive recordings (SEEGformer) or high-density configurations (DeepSOZ, MBAT), leaving a gap in generalisability to the routine 18-channel low-density montage that constitutes the vast majority of clinical practice.

Beyond these domain-specific models, Transformer architectures have been widely adopted in broader EEG epileptic analysis, though predominantly for detection rather than localisation. For seizure detection, hybrid LSTM-Transformer models have achieved 99.75% sensitivity on CHB-MIT [28]; a multidimensional Transformer with RNN fusion has also been proposed for seizure classification [29]. For interictal epileptiform discharge (IED) detection, Transformer-based classifiers have achieved multi-label classification accuracies of 93.47% on paediatric scalp EEG [30], dissociated physiological ripples from epileptiform discharges [31], and achieved 99.8% accuracy in patients with self-limited epilepsy with centrotemporal spikes [33]. For seizure subtype classification, multi-branch mutual-distillation Transformers have addressed cross-subject generalisation on small, labelled datasets [32]. More broadly, Transformers capture temporal information in EEG more effectively than CNNs or RNNs, with greater focus on global features of brain functional activity [34]. However, most existing EEG Transformers flatten temporal and channel dimensions into a single sequence, treating temporally adjacent samples and spatially adjacent channels as positions in a homogeneous sequence.

In summary, these challenges stem from methodology gap: existing deep learning frameworks treat non-invasive scalp EEG as if it were high-SNR intracranial data, failing to account for the physical constraints of volume conduction and the multi-scale nature of interictal discharges. This misalignment creates a vicious cycle where models perform well in narrow laboratory settings but fail in clinical reality, ultimately eroding clinician trust. To bridge this gap, an effective architecture must be designed from the outset specifically for large-cohort, low-density scalp EEG to achieve robust SOZ localisation. This requires overcoming two explicit limitations of current generic models: first, their fixed receptive fields cannot capture the wide temporal range (20–200 ms) of multi-scale interictal events; second, standard Transformers mathematically conflate temporal phase shifts with spatial electrode displacements, losing crucial physiological context. Our proposed framework directly addresses these architectural bottlenecks by decoupling spatial and temporal configurations within a low-density clinical montage.

### 1.4 Study Rationale and Objectives

The preceding review of the literature (Section 1.3) identifies two critical gaps in current deep learning frameworks for SOZ localisation: first, standard Transformers flatten time and channels into a single sequence, confounding temporal phase shifts with spatial propagation topographies, fail to counter volume conduction; second, most high-performing models are validated on small, invasively recorded cohorts, leaving a substantial gap in generalisability to routine low-density scalp EEG.

Overall objective: This study aimed to demonstrate proof-of-principle that a deep learning framework can reliably localise the SOZ at the lobar level from routine low-density scalp EEG. We suggest that design features specifically address the physical limits of scalp recordings, i.e. factorised spatiotemporal reasoning to disentangle waveform evolution from spatial propagation, and clinically grounded anatomical priors can provide clinically useful results without the need for invasive recordings or dense electrode arrays.

To test this hypothesis and demonstrate translational viability, we pursue five interlocking strategies that collectively address the generalisability, robustness and interpretability requirements of a deployable clinical tool.

Firstly, build a diversified and well-annotated training set. We systematically screen and annotate a large cohort of 100 unilateral focal epilepsy patients (262 sessions, >6,000 hours) from the Harvard EEG Database, with ground-truth lobar labels obtained from expert clinical reports. This cohort is highly heterogeneous in terms of age (22-85 years) and balanced hemispheric lateralisation (56% left, 44% right) which provides the necessary heterogeneity of data to support cross-patient generalisation.

Secondly, design and ablate a physiologically motivated architecture. We present SOZNet, a factorised spatiotemporal Transformer that explicitly isolates temporal and spatial attention pathways, a direct architectural solution to volume conduction that standard Transformers confound by flattening time and channels into a single sequence. We augment this with a hemispheric mutual exclusivity penalty directly encoded in the loss function that enforces the clinical axiom of unilateral focal epilepsy during training. Systematic ablations quantify the contribution of each component.

Thirdly, apply data augmentation in a targeted manner. We employ an online augmentation pipeline to avoid overfitting and impose invariance to inter-patient and inter-session variability, especially electrode impedance fluctuations, EMG artefacts, and placement variations, including Gaussian noise injection, amplitude scaling, random temporal cropping, and hemispheric channel swap. These augmentations increase the effective diversity of the training samples without label leakage.

Fourthly, Test in clinically realistic generalisation scenarios. Unlike most studies that evaluate on held-out data from the same institution and recording protocol, we perform a stress test of our framework on three progressively more difficult scenarios: (i) an internal holdout set from the same institution, but with part of patients without ictal annotations, testing reliance on interictal patterns alone; (ii) a cross-institutional external set from Boston Children’s Hospital, with different equipment, electrode placement protocols, and a pediatric demographic; (iii) wholly unlabelled random EEG segments, where no spike timestamps are available, mimicking the reality of routine clinical recordings that lack dense manual annotations.

Fifthly, establish mechanistic interpretability and safety. Clinical adoption demands more than aggregate metrics. We therefore conduct: (a) case-level trajectory analysis to show that predictions track with expert summaries across continuous recordings; (b) zero-false-positive verification on healthy control recordings.

This framework provide a non-invasive, efficient, and clear method to speed up presurgical evaluation. It could reduce the need for long hospital stays, medication tapering, and invasive electrode placement. At the same time, it would offer automated decision support to help plan stereotactic EEG implantation. Crucially, our model is trained to predict the SOZ as defined by expert clinical EEG interpretation; its output is a decision-support tool to inform, rather than replace, the epileptologist’s judgement of the EZ.

## 2: Methodology

### 2.0 Methodological Rationale: A Three-Stage Pipeline

Before detailing the implementation, we outline the high-level logic governing our methodological choices. The pipeline operates in three conceptual stages, each directly addressing a physical constraint identified in Section 1.3:

1. **Signal purification (Sections 2.1–2.5)**: Raw scalp EEG is contaminated by muscle artefacts, line noise, and baseline drift. We apply a standardised preprocessing cascade - bipolar re-referencing, wavelet denoising, and common-average referencing to isolate genuine neural signals while preserving spike morphology.
2. **Multi-scale feature extraction (Section 2.9.1–2.9.2)**: Interictal spikes have broad durations (20–200 ms). Instead of trying to find the one optimal size of the filter, our front-end employs four parallel convolutions with different kernel lengths, analogous to a pathologist examining tissue at multiple magnifications to capture both sharp transients and slow after-waves.
3. **Factorised spatiotemporal reasoning (Section 2.9.3–2.9.5)**: Volume conduction spreads one cortical source across several electrodes. Standard Transformers mix up the “when” of a signal with the “where” of a signal by collapsing time and channels into one sequence. We explicitly decompose these two dimensions into independent attention pathways as our key innovation, which are only combined at the decision layer. This requires the model to learn the evolution of the waveform and the spatial propagation as separate physical events, as clinicians do instinctively when reading EEG.

Figure 3 shows the full pipeline overview of the experiment step by step:

**Figure 3:**
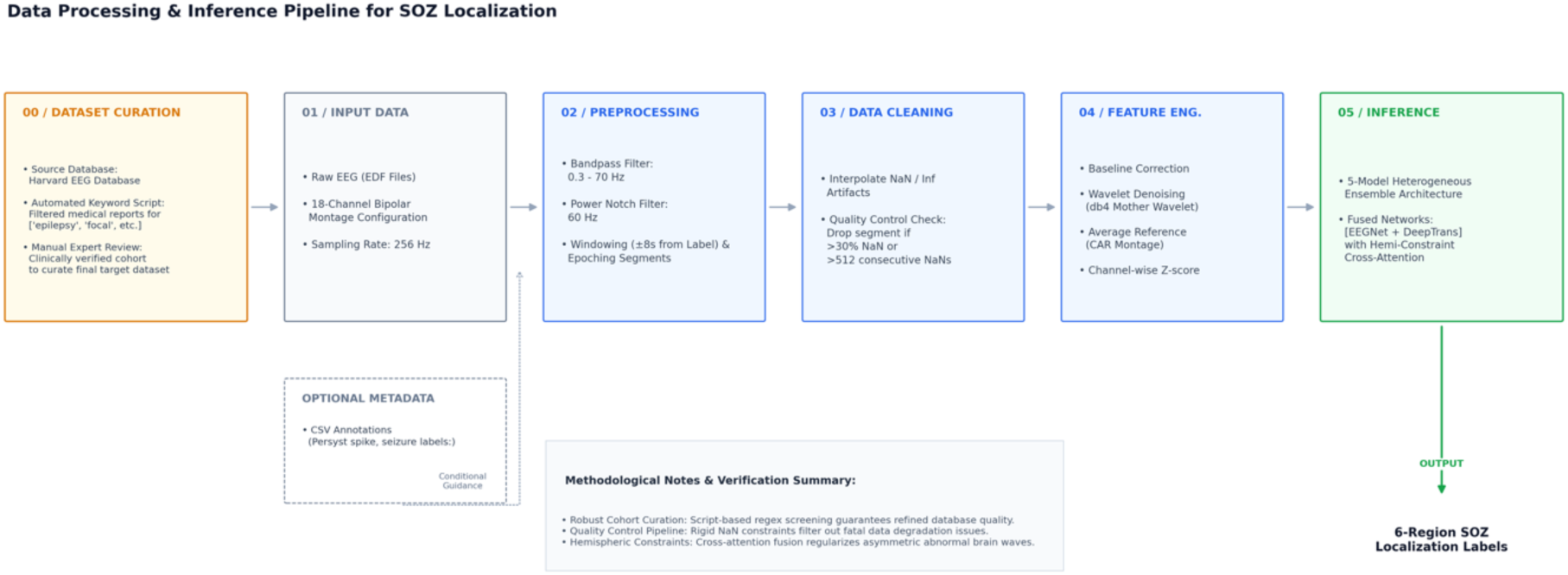
End-to-end data processing and inference pipeline for six-region SOZ localisation. The computational workflow comprises six sequential phases: (00) Dataset curation incorporating clinical expert validation and multi-centre test partitioning; (01) Raw 256 Hz scalp EEG ingestion mapped onto a standard 18-channel longitudinal bipolar configuration; (02) Preprocessing via zero-phase bandpass/notch filters and deterministic temporal windowing; (03) Data cleaning leveraging automated linear interpolation and strict signal quality thresholding (max consecutive NaNs = 512, max NaN ratio = 0.3); (04) Feature engineering executing multi-scale wavelet denoising (db4, VisuShrink) and spatial re-referencing (Common Average Reference); and (05) Inference driven by a heterogeneous multi-model ensemble neural architecture (M1–M3, fused via model soup and soft voting). The pipeline ultimately yields robust multi-label probability distributions indicating localisation across six anatomical cortical regions.

### 2.1 Data acquisition and case selection

All data utilised in this study were obtained from the Harvard Electroencephalography Database version 4.1 (HEEDB) [35], which aggregates clinical recordings from four medical centres: Massachusetts General Hospital (MGH), Brigham and Women’s Hospital (BWH), Beth Israel Deaconess Medical Centre (BIDMC), and Boston Children’s Hospital (BCH).

While MGH and BWH provide more raw EEG files, many are short outpatient recordings with higher noise levels and sparse labels. These are less suitable for training deep learning models. In contrast, the BIDMC cohort predominantly features 24-hour continuous EEG (cEEG) recordings, providing prolonged observation windows and a higher density of localised spike and seizure labels that are essential for model training. This study focuses on the BIDMC cohort. This single-site selection, while ensuring annotation density and recording standardisation, may introduce site-specific biases in patient demographics and referral patterns; the external validation on the BCH cohort is designed to provide preliminary evidence of cross-institutional generalisability.

To select the target clinical cohorts, we implemented a two-stage screening process consisting of automated lexical scoring followed by manual expert-report verification.

#### Stage 1: Weighted Rule-Based Text Screening

A customised natural language processing (NLP) script was used to parse the official clinical reports authored by epileptologists. Patient sessions were quantitatively evaluated using a weighted rule-based grading matrix:

- High-Relevance Indicators (+2 points): Explicit positive descriptors signalling localised or active ictal events, such as “focal”, “electrographic seizure”, and “clinical seizure”, were assigned a weight of +2 per occurrence.
- Moderate-Relevance Indicators (+1 point): General epileptiform findings, including “discharge”, were assigned +1 point.
- Exclusionary Modifiers (-1 or -2 points): Negative qualifiers demonstrating the absence of relevant activity (e.g., “no electrographic seizure”) resulted in a deduction of 1 or 2 points to mitigate false positives from keyword matching.

Sessions achieving scores above a predefined threshold (2) were selected for the next stage, 3874 reports passed the lexical selection.

#### Stage 2: Manual Report Verification and Exclusion Criteria

Following lexical scoring, a manual audit of the candidate clinical reports was conducted by reviewing the expert clinical summary for each session. For each report, we extracted:

1. Seizure onset side: documented as left, right, or bilateral (bilateral excluded)
2. Seizure onset lobe: documented as frontal, temporal, parietal, occipital, or combinations (e.g., temporo-parietal)
3. Semiology concordance: whether clinical seizure semiology matched the electrographic onset.

Based on these clinical descriptions, each session was assigned a six-region multi-label ground-truth vector according to the anatomical mapping defined in Section 1.1. For example, a report stating “seizure onset in the left temporal lobe with propagation to the left parietal region” was labelled as [0, 0, 1, 0, 0, 0] - where the ‘1’ indicates the left temporo-parietal region (R3). Sessions with documented bilateral onset, ambiguous localisation, or lacking a clear SOZ in the final report were excluded for this proof-of-principle study.

### 2.2 Ethics and Data Access

All data utilised in this study were obtained from the Harvard Electroencephalography Database version 4.1 (HEEDB), a publicly accessible repository that aggregates de-identified clinical EEG recordings from four partner institutions: Massachusetts General Hospital (MGH), Brigham and Women’s Hospital (BWH), Beth Israel Deaconess Medical Centre (BIDMC), and Boston Children’s Hospital (BCH). The HEEDB dataset was generated through retrospective aggregation of routine clinical recordings, with all personally identifiable information removed before release. Specifically, patient names, medical record numbers, dates of birth (converted to age at recording), and all free-text clinical notes containing potential identifiers were stripped through an automated de-identification pipeline. Only non-identifiable demographic information (age, sex), recording metadata (duration, electrode montage), and the clinical reports (with identifiers redacted) are retained in the distributed dataset. The dataset was pre-approved for secondary research by the IRB of each contributing institution. Details are documented in the HEEDB release paper.

Access to the HEEDB v4.1 dataset was granted through a formal data use agreement with the repository administrators. We completed the required data access training, covering (i) responsible handling of protected health information (PHI) as defined under HIPAA, (ii) data storage and encryption requirements, and (iii) restrictions on data redistribution and re-identification attempts. Access was provided via secure, password-protected institutional servers with role-based authentication, for the duration of the research project (Dec 2025-Aug 2026). No identifiable patient data were stored on local workstations; all analyses were conducted on institutionally managed servers.

Because the dataset is fully de-identified and anonymised before distribution, this study does not meet the regulatory definition of human subjects’ research requiring individual informed consent. The residual ethical risks are minimal and primarily relate to: (1) the potential for indirect re-identification through rare phenotype combinations (e.g., age, sex, and rare epilepsy subtype), which is mitigated by the large cohort size (100 patients) and the aggregation of recordings across multiple centres; and (2) the general risk of data breaches, which is addressed through the institutional data governance framework and secure server infrastructure. No patient data were accessed outside the approved research context, and all analyses were conducted in accordance with the ethical principles outlined in the Declaration of Helsinki. Individual informed consent was not required for this secondary analysis, as the HEEDB dataset was fully de-identified before distribution and approved for secondary research use by the IRB of each contributing institution, consistent with HIPAA guidelines for de-identified data (45 CFR 46.104).

### 2.3 EEG Data Harmonisation, Window Cropping, and Quality Control Pipeline

#### 2.3.1: Channel Standardisation

EEG recordings vary in channel layout. We standardised all recordings by: (1) removing non-EEG channels (e.g., oxygen saturation, pulse rate); and (2) applying a consistent spatial referencing scheme.

We applied band-pass filtering (0.3–70 Hz) to remove slow drifts and high-frequency noise, plus a notch filter at 60 Hz to remove electrical interference from power lines [36].

#### 2.3.2: Multi-Format Annotation Alignment and Time-Locked Epoching

Automated epileptiform labels (spikes and seizures) were parsed from CSV files generated by the Persyst software. There was a small time offset between the EEG signal and the Persyst annotation file. We corrected this by aligning both to the “start recording” marker.

To remove non-epileptiform behavioural or environmental markers logged during the session, a vocabulary exclusion filter was applied. Irrelevant textual descriptions, such as ocular movements (“eyes open/closed”), physiological artefacts (“chewing”, “movement”), and operational event logs (“montage review”, “video fault”), were removed to isolate objective spike and seizure annotations. Using these spike and seizure markers as temporal anchors (t=0), the continuous streams were segmented into 16-second windows (from −8.0 seconds to +8.0 seconds).

We did not perform manual verification of individual windows, as this was infeasible at the scale of our dataset; instead, use an NLP filter to exclude artefact-contaminated segments, reducing false positives. Moreover, our spatial ground-truth labels derive solely from the final expert clinical reports, not from Persyst confidence scores.

#### 2.3.3: Dimension Standardisation, Resampling, and Parallel Serialisation

Recordings had different sampling rates (256 Hz, 512 Hz, or 1024 Hz). We resampled all windows to 256 Hz, giving each window 4,096 time steps (16 seconds × 256 Hz). Spatial or temporal boundary variations were resolved via centralised trimming or symmetrical zero-padding, ensuring a uniform tensor shape of 18 × 4096 across all windows. The standardised segments were serialised into binary PyTorch tensors. Implementation details of the parallel processing pipeline are provided in Appendix B.

### 2.4 Data Cleaning: Signal Preprocessing and Quality Control

To ensure signal integrity and avoid mathematical instabilities during training, an automated data-cleaning and quality-control protocol was applied to all extracted EEG segments (standardised shape 1 × 18 × 4096).

Infinite values were converted to NaN. A two-tiered quality control protocol was applied: segments exceeding either a maximum consecutive NaN threshold or a global missing ratio threshold were discarded. For remaining segments, missing values were imputed per channel using 1D linear interpolation, with nearest-neighbour extrapolation at sequence boundaries. The full set of rejection thresholds and interpolation parameters is provided in Appendix B.

### 2.5: Signal Enhancement and Feature Normalisation

Following serialisation and quality control, the verified .pt segments underwent a digital signal processing (DSP) pipeline to remove transient high-frequency noise and normalise amplitude scales before model training.

1. Temporal Baseline Correction: We applied temporal baseline correction to mitigate low-frequency drift and baseline wander caused by slow movement or electrode-tissue polarisation. The initial 100-time steps of each segment were designated as the reference window, and its localised channel-wise mean was subtracted from the entire sequence to realign the signal’s electrical baseline to zero [37].
2. To suppress high-frequency muscle artefacts and line noise without attenuating neural spikes, a discrete wavelet transform denoiser was applied. Each channel was decomposed into four levels using the Daubechies 4 (db4) mother wavelet, with adaptive universal thresholding (VisuShrink) [38]. Soft-thresholding was applied to detail coefficients across all scales, and the signal was reconstructed via inverse DWT. Detailed parameter settings are provided in Appendix C.
3. Spatial Re-referencing (Common Average Reference): To eliminate residual environmental common-mode noise remaining after the initial bipolar montage, a Common Average Reference (CAR) filter was applied spatially. The instantaneous mean voltage across all 18 channels was calculated and subtracted from each channel.[38]
4. Z-Score Standardisation and Incremental Logging: Finally, to optimise numerical stability and ensure stable gradient flow during training, channel-wise Z-score standardisation was executed, scaling each channel sequence to possess zero mean and unit variance.

### 2.6 Dataset Cohort and Label Distribution Analysis

The study cohort comprises 100 focal epilepsy patients with 262 continuous video-scalp EEG recording sessions, totalling over 6,000 hours (original file length). To evaluate generalisability across different clinical demographics, we divided the data into three splits: a primary training/internal set (BIDMC, N=80 patients for training, N=10 for validation, N=10 for internal test), a held-out set from the same institution (BIDMC, N=28 patients,

annotation-sparse), and an external cross-institutional test set (Boston Children’s Hospital, N=9 patients)(Figure 4.a-c).

**Figure 4:**
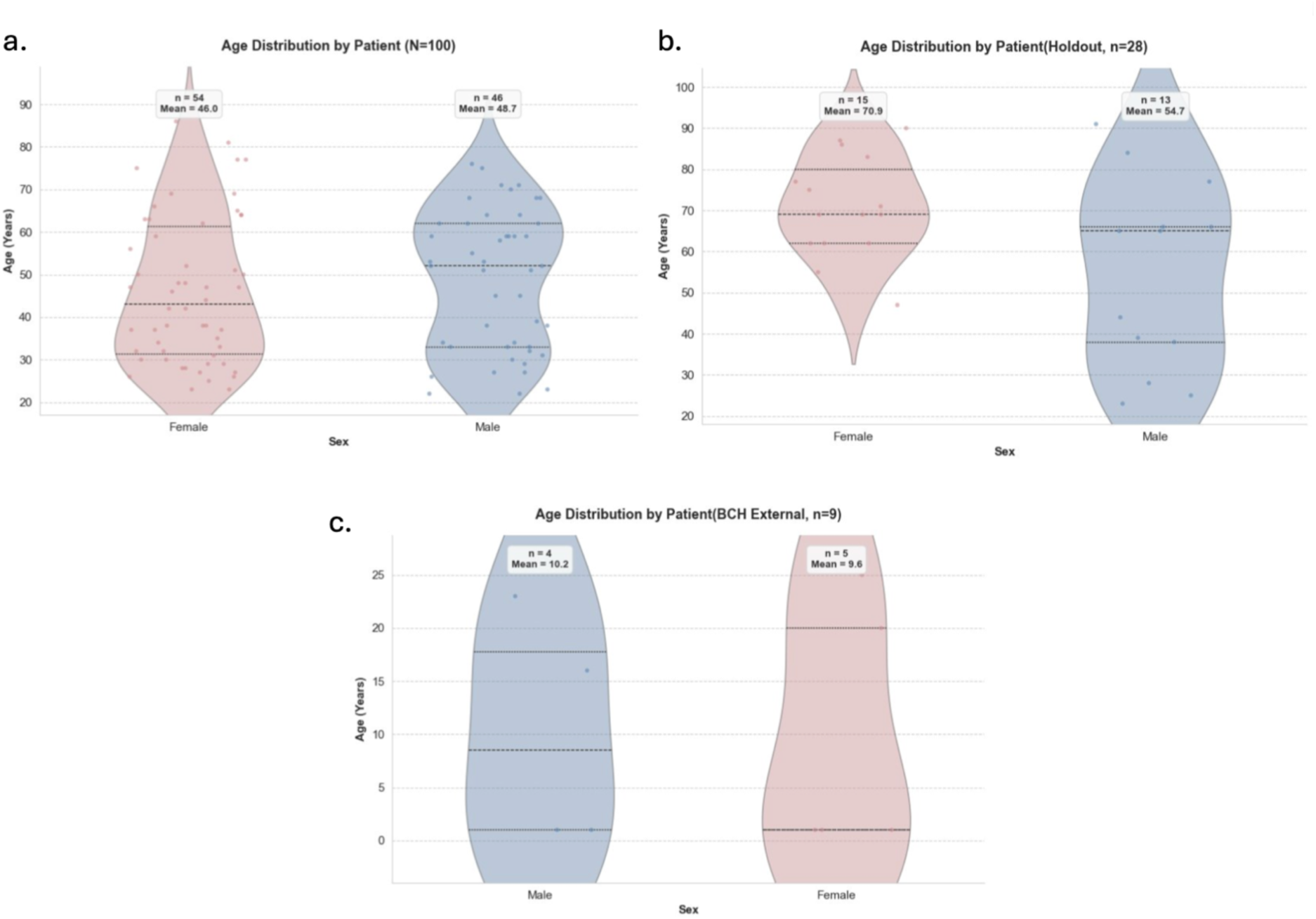
Demographic characteristics of the study cohort. (a) Violin plots showing age distribution by sex across 100 patients. The male subgroup (n = 46) has a mean age of 48.7 years (range: 22–76); the female subgroup (n = 54) has a mean age of 46 years (range: 23–86). The female cohort exhibits a broader age distribution and larger sample size. (b) showing age distribution by sex on the holdout set with 28 patients (Male=13, Female=15), with a male mean age of 54.7 and 70.9 for females. Lower panel: 9 patients from BCH, 5 of them are paediatric cases (age ≤ 1).

Demographic profiles across cohorts. We compared the age and sex distributions across these three cohorts (Figure 4). The primary BIDMC cohort exhibits a symmetrical, spindle-shaped age distribution. The male subgroup (n=46) has a mean age of 48.7 (range: 22–76), while the female subgroup (n=56) has a mean age of 46 years (range: 23–86), reflecting a broader age range. In contrast, the holdout cohort (n=28) is much older, with a female mean age of 70.9 and a male mean of 54.7, representing an older-adult demographic. The external BCH cohort (n=9) shifts toward a paediatric/adolescent demographic (n=5 are less than 1 year old). This demographic heterogeneity across training, held-out, and external sets provides a stress test for the model’s robustness to age- and sex-related EEG variability, as paediatric and geriatric EEGs exhibit distinct background rhythms, faster/slower dominant frequencies, and different morphological characteristics of interictal discharges compared to the middle-aged adult population.

Patient-level label distribution. Under our multi-label clinical framework, each patient is mapped to one or multiple anatomical target regions reflecting their epileptogenic network. To avoid the statistical inflation caused by repeated session-level counts (where some patients contribute many long recordings), we aggregated the ground-truth labels at the patient level (N=100 unique patients with definitive unilateral SOZ) (Figure 5).

**Figure 5:**
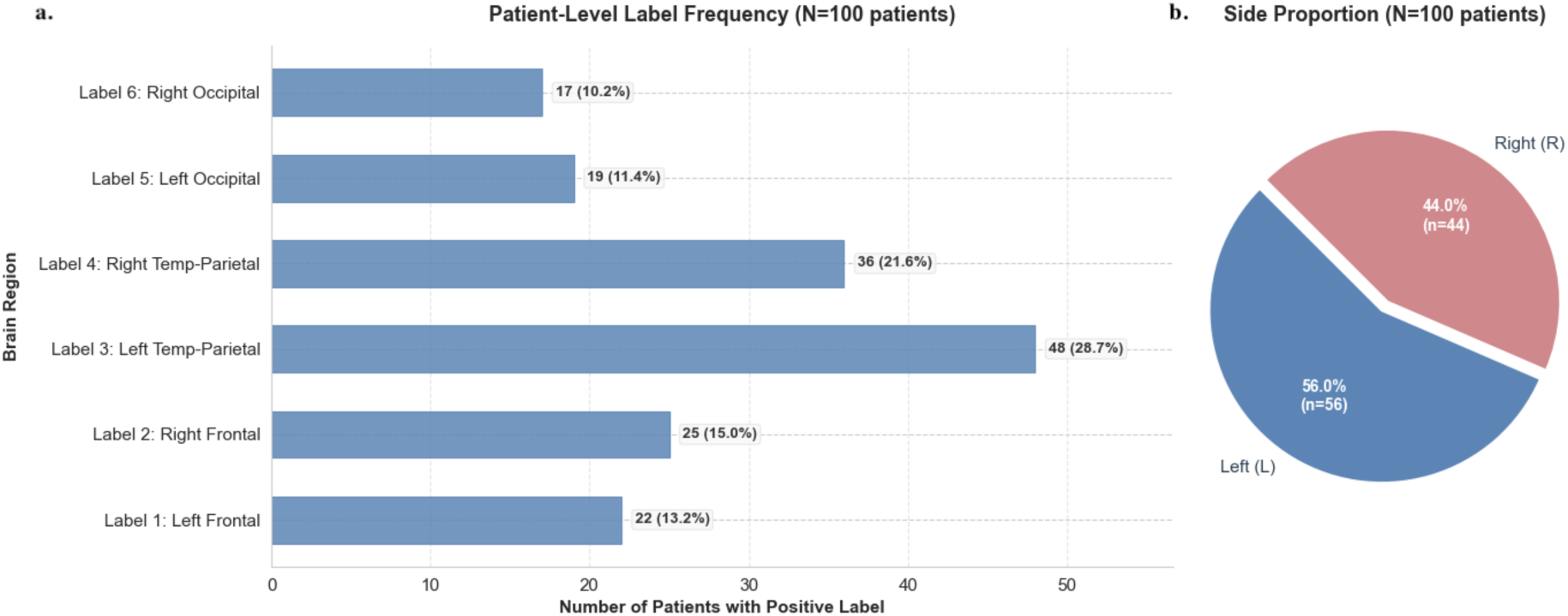
Distribution of SOZ labels across six anatomical regions. (a) Absolute counts and percentages of positive labels across the six regions. Left Temporo-Parietal 28.7% and Right Temporo-Parietal 21.6% constitute most cases, consistent with the epidemiological predominance of temporal lobe epilepsy (TLE) and its propagation to adjacent parietal cortex. The remaining four regions have pretty much identical distribution (10.2-15%). (b) shows the percentage of overall lateralisation of the dataset ( L=56%, R=44%)

The label distribution matches clinical epidemiology: temporo-parietal regions are the most common, accounting for 28.7% (left) and 21.6% (right) of patients. This is consistent with the well-known predominance of temporal lobe epilepsy, which accounts for over 60% of focal epilepsy cases [39]. Furthermore, from an electroanatomic perspective, the temporal-parietal neocortex covers a significantly larger volume and surface area under the standard 10-20 system compared to the occipital pole [8]. The remaining 4 regions show moderate representation: Left Frontal: 13.2%; Right Frontal: 15.0%; Left Occipital: 11.4%; Right Occipital: 10.2%.

To evaluate cross-patient generalisability, we designated two independent test cohorts: a holdout set from the same institution (87 sessions, 28 patients) and an external set from Boston Children’s Hospital (13 sessions, 9 patients). The holdout includes annotation-sparse sessions without ictal events; the BCH cohort provides cross-institutional validation with different recording equipment and protocols.

Hemisphere-level lateralisation shows a left-sided predominance (56% Left vs. 44% Right hemisphere), confirming a well-balanced unilateral distribution without extreme hemispheric bias.

### 2.7 Cohort Partitioning and Patient-Level Data Splitting

The dataset (100 patients, 262 sessions) was split at the patient level using an 8:1:1 ratio. This yielded 80 patients for training, 10 for validation, and 10 for independent testing. Crucially, all sessions from a single patient were assigned exclusively to one subset, preventing data leakage and spatial co-adaptation. This guarantees that the model’s performance reflects true cross-subject generalizability to unseen neural anatomies rather than memorisation of individual traits.

#### Data Augmentation & Model Architecture, Design

##### 2.8.1 Data Augmentation and Spatiotemporal Perturbation Strategy

From the 80-patient training cohort, approximately 55,000 non-overlapping 16-second windows were extracted using the spike- and seizure-anchored protocol described previously. To mitigate overfitting and enforce invariance against inter-patient anatomical and impedance variability, we apply a stochastic online augmentation pipeline (detailed below) to each standardised 1 × 18 × 4096 tensor, yielding an extra 50,000 copies of 16-second chunks from data augmentations in every training epoch. The following perturbations are applied dynamically during training (illustrated in Figure 6):

**Figure 6:**
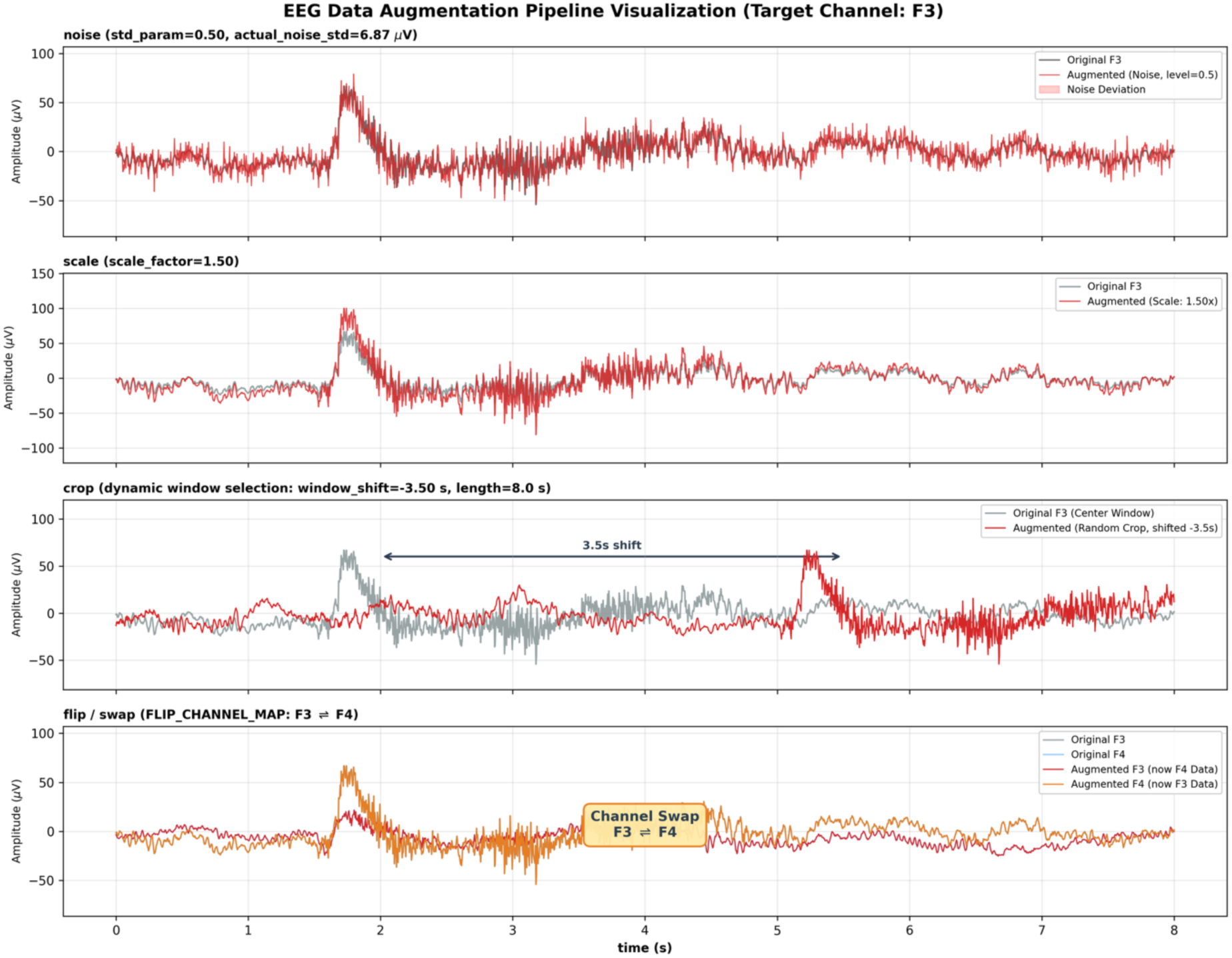
Visualisation of online data augmentation strategies applied during training. Each panel shows a real training window from the F3 channel (18 × 2048 time steps). Top-left: Original signal (baseline). Top-right: Gaussian noise injection (σ∼U(0.1,0.5)) simulating EMG and thermal circuit noise. Bottom-left: Random amplitude scaling (α∼U(0.8,1.2)) simulating electrode-skin impedance fluctuations, forcing the model to rely on relative spike morphology rather than absolute voltage magnitudes. Bottom-right: Dynamic temporal cropping with random shift (δ∼U(−3.5,+3.5) s) preventing overfitting to a fixed pre-ictal baseline phase. Hemispheric channel swap (not shown) is applied as a geometric regulariser with 0.5 probability. All augmentations are applied online during training, yielding 50,000 augmented 8-second chunks per epoch from 55,000 original 16-second windows.

We apply four online augmentations (Figure 6) to enforce invariance and reduce overfitting:

1. Gaussian noise σ ∼ U(0.1, 0.5) to simulate EMG/thermal noise;
2. Amplitude scaling (0.8,1.2) to simulate impedance fluctuations, making the model rely on morphology over absolute voltage;
3. Random temporal croppingδ∼U (−3.5, +3.5) s to avoid overfitting to a fixed pre-ictal baseline;
4. Full hemispheric channel swap (left → right, right → left) as a geometric regulariser against electrode placement variability. We evaluate its impact in Section 3.4.

##### 2.8.2 Session-Level Sampling Strategy

To prevent overfitting to individual long-duration sessions, we cap the maximum number of windows extracted from any single session at Nmax = 300. This ensures that a single patient with much more spike labelling doesn’t dominate the training distribution.

For each of the three SOZNet seeds and the two ablation variants (detailed in Appendix D), we apply different random subsampling trajectories to the training windows. This strategy increases the effective sample diversity seen by the ensemble; each model sees a slightly different “view” of the training data without artificially inflating the dataset size or introducing label leakage. This strategy, combined with the online augmentations above, further mitigates the risk of session-specific overfitting.

### 2.9 Spatiotemporal Neural Architecture: From Physiology to Implementation

The overall architecture of the proposed SOZNet is illustrated in Figure 7. It comprises three stages: a multi-scale convolutional front-end that captures heterogeneous discharge morphologies, a depthwise spatiotemporal convolution that fuses cross-channel information, and a factorised Transformer that decouples temporal and spatial attention into independent pathways, a direct architectural response to the volume conduction constraints discussed in Section 1.3.

**Figure 7:**
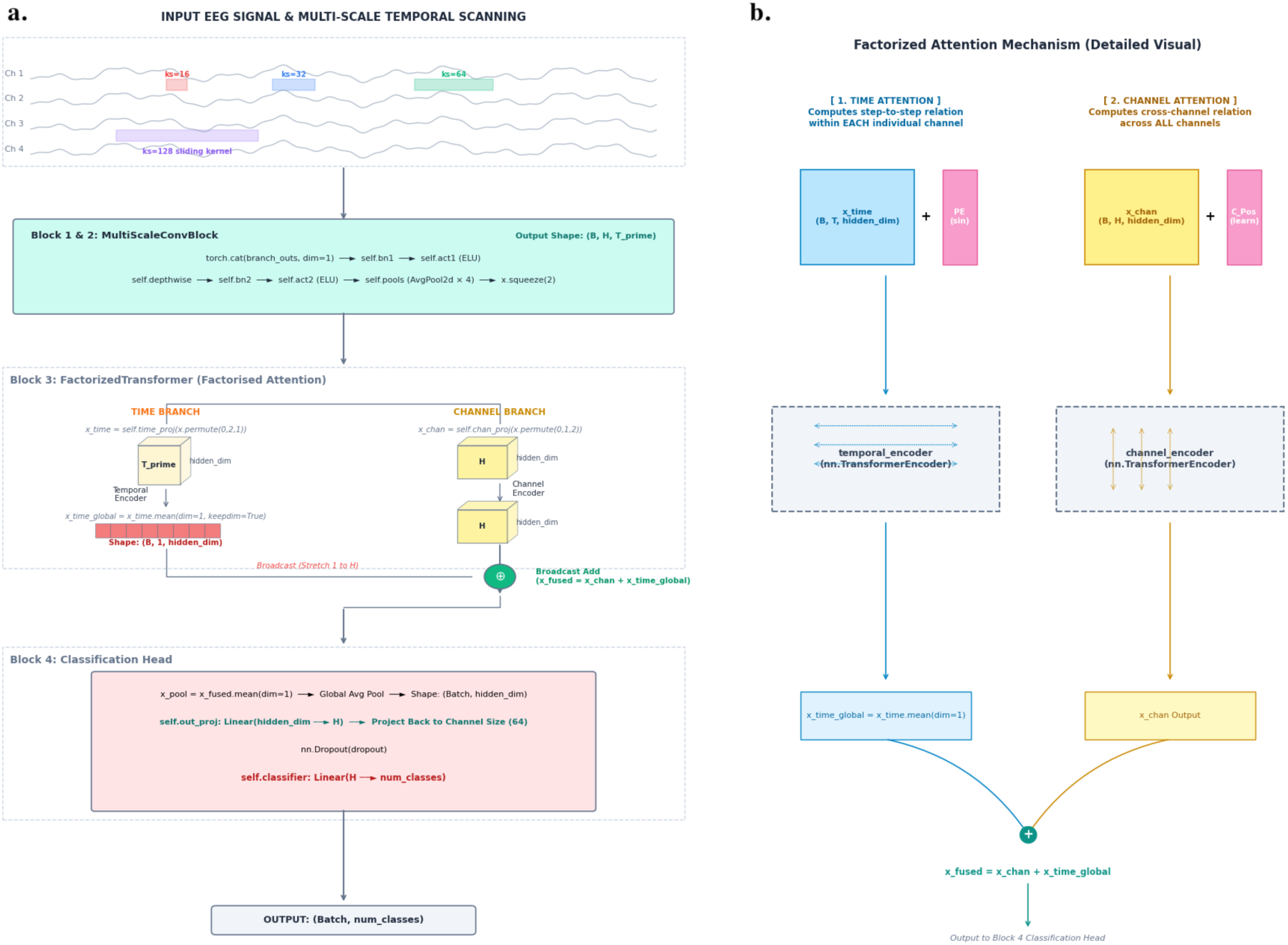
M1 architecture (SOZNet) with factorised spatiotemporal Transformer. (a) Overall architecture comprising three stages: (i) a multi-scale convolutional front-end (MultiScaleConvBlock) with parallel kernel sizes {16, 32, 64, 128} capturing heterogeneous discharge morphologies (20–70 ms spikes, 70–200 ms sharp waves, 4–12 Hz slow waves); (ii) depthwise spatiotemporal convolution with feature concatenation and temporal downsampling (×16), producing a spatiotemporal token tensor X∈R ^B×64×128^; and (iii) a Factorised Transformer (see b panel) that decouples temporal and spatial attention. (b) Factorised Transformer showing: (i) dual-path projective embedding (temporal: B×128×128; spatial: B×64×128); (ii) asynchronous positional encodings (sinusoidal for temporal, learnable for spatial); (iii) factorised attention encoders (2 layers, 4 heads, hidden dimension 128); and (iv) low-rank consensus fusion where global temporal pooling (B×1×128) is broadcast and added to spatial features, producing fused representation X fused ∈ R ^B×64×128^. The fused tensor is pooled and passed to a linear classifier producing six sigmoid logits.

**Figure 8:**
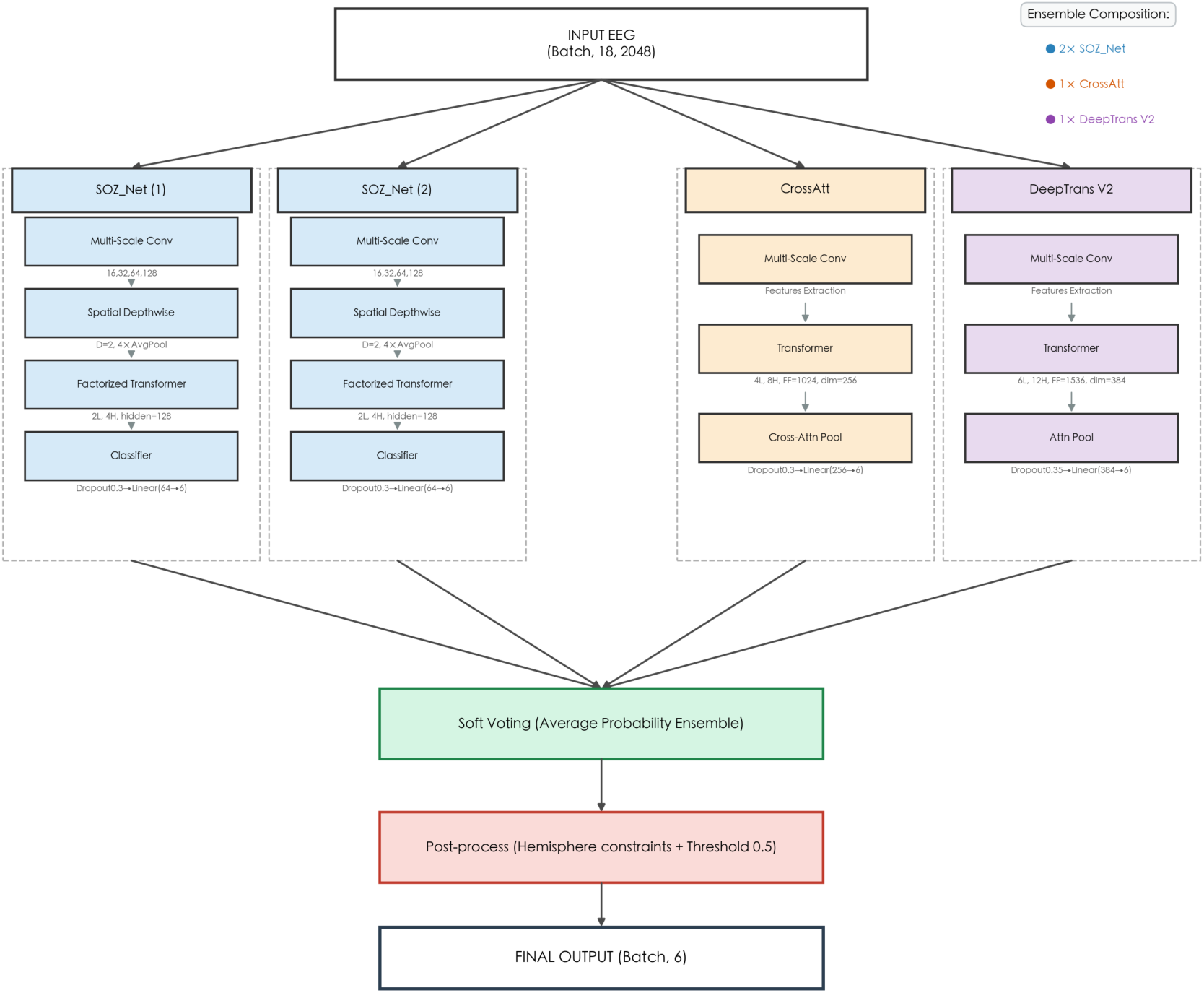
Heterogeneous ensemble strategy with model soup and soft voting. The final ensemble comprises five constituent models: three M1 variants (seeds 100, 101, 102), one M2 variant (seed 42), and one M3 variant (seed 43). Stage 1 (Model Soup): For each architecture, three independent fine-tuned variants (differing in Gaussian noise injection levels) are averaged element-wise to produce a fused model: 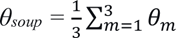. Stage (Soft Voting): At inference time, each fused model produces a six-dimensional probability vector p^(m)^=σ(z^(m)^); the ensemble prediction is the arithmetic mean 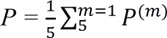. This hierarchical integration balances intra-architecture consolidation with inter-architecture diversity, reducing both bias and variance without incurring inference-time ensemble overhead.

#### 2.9.1 Physiological Motivation: Capturing Spikes from 20 ms to 200 ms

Interictal events occur at very different time scales: brief spikes (20–70 ms), longer sharp waves (70–200 ms), and slow-wave rhythms (4–12 Hz, lasting hundreds of milliseconds). A single fixed-size detector cannot capture all of these. Our MultiScaleConvBlock uses four parallel convolutions with different kernel sizes to detect events at all these scales simultaneously.

We introduce a parallelised multi-receptive-field front-end, the MultiScale Temporal Scanning Block (MultiScaleConvBlock). This module processes the input through four independent 1D-convolutional pathways with distinct kernel sizes:

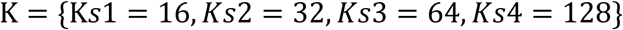

At 256 Hz sampling rate, kernel sizes of 16 and 32 correspond to corresponding to receptive fields of 62.5 ms and 125 ms, which cover the 20–70 ms duration of brief spikes. The larger kernels (64 and 128) extend to 250 ms and 500 ms, encompassing the longer sharp-wave and slow-wave complexes (70–200 ms) that frequently follow interictal spikes. The micro-scale branches (ks=16,32) are designed for sharp spike contours, while the macro-scale branches (ks=64,128) target sustained slow-wave rhythms. These branches jointly capture multi-scale temporal features for downstream modules.

#### 2.9.2 Spatial Fusion via Depthwise Convolution: Depthwise Convolution

Let the input segment be denoted as X ∈ R ^B^ ^×1^ ^×C^ ^×T^, where B signifies the batch size, C = 18 represents the number of bipolar channels (num_electrodes), and T indicates the discrete temporal sequence length (chunk_size). Within the MultiScaleConvBlock, the input tensor passes through a bank of parallel 2D convolutions (nn.ModuleList)with kernels constrained to (1, ks) for k ∈ {16, 32, 64, 128}. This restricts early feature extraction to channel-isolated temporal filtering, preserving the spatial dimension for later fusion. Symmetric padding of k//2 is applied to maintain temporal length across branches.

Each branch produces an intermediate feature map of depth F1=8. These are concatenated along the channel dimension:

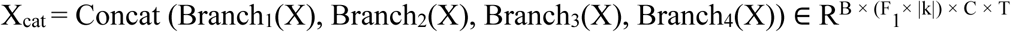

followed by Group Batch Normalisation and ELU activation.

To fuse cross-channel spatial information without the quadratic parameter growth of standard 2D convolutions, we apply a depthwise spatial convolution (self.depthwise). This operator uses a kernel of size (*C*,1) with groups equal to the input feature depth F1×∣K∣=32, thus pooling across all 18 spatial channels independently per feature layer. An output multiplier D=2 yields an amplified depth:

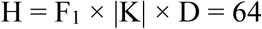

A second Batch Normalisation and ELU activation follow. Subsequently, the tensor is passed through four sequential 2D average-pooling layers with kernel (1,2) and stride (1,2), compressing the temporal dimension by a factor of 2^4^: T to T’ = T // 16 (T’_prime = 128). (Given the original T=2048 after chunking). Finally, x.squeeze(2) removes the singleton spatial dimension, producing the regularised spatiotemporal token tensor:

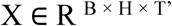

#### 2.9.3 Architecture Rationale: Why Separate Time and Space?

Unlike invasive sEEG with high SNR, scalp EEG suffers from severe volume conduction and spatial smearing, where a single cortical source projects diffusely across multiple channels. In standard Transformers, flattening time (T) and space (H) into a single sequence forces the attention matrix to model cross-products between temporally distant samples and spatially distant channels simultaneously. This is computationally prohibitive, O((H × T)^2^), and conflates temporal phase shifts with spatial topography. That is, the model cannot distinguish whether a signal arrived 20 ms later or originated from a different electrode, yet these scenarios have entirely different clinical implications. Our factorisation explicitly disentangles these two physical entities, allowing temporal attention to model waveform evolution and spatial attention to model inter-channel propagation independently

To enforce strict geometric factorisation, the proposed network splits the tensor Xout into two parallel, decoupled embedding paths using distinct linear projection layers (self.time_proj and self.chan_proj). Spatial Channel Path Projection (self.chan_proj): Concurrently, the tensor is processed along its spatial coordinate, treating the H channels as the sequence length. A linear layer projects the downsampled temporal steps T’ into the same latent dimension:

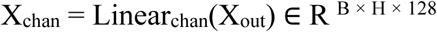

This dual-path projective embedding maps the raw data into two symmetrical matrices, allowing the downstream encoders to process pure, unmixed temporal dynamics and pure spatial topology independently. This projection maps the raw EEG into two separate representations: one encoding temporal waveform evolution, the other encoding spatial electrode topography.

#### 2.9.4 Factorised Attention: Implementation Details (Asynchronous Encoding & Dual Encoders)

We inject asynchronous positional encodings tailored to each branch. Because time and space have fundamentally different physical meanings in EEG, time is sequential and causal, while space is discrete and topological, we introduce an Asynchronous Positional Encoding mechanism customised for each branch:

Sinusoidal Temporal Encoding (self.temporal_pos): The temporal path contains strict sequential constraints (i.e., the chronosynchronous flow of the spike waveform). Thus, a deterministic, non-trainable sinusoidal positional matrix Psin ∈ R ^1^ ^×^ ^max_len^ ^×^ ^128^ is added to Xtime:

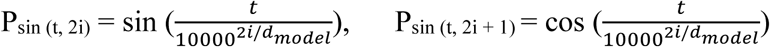

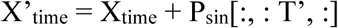

Learnable Spatial Channel Encoding (self.channel_pos): Conversely, the spatial pathway maps discrete channels across the 10–20 grid. Since these channels lack a natural linear order, we use a fully trainable parameter tensor Pchan ∈ R ^1^ ^×^ ^1^ ^×^ ^128^ is initialised and added via broadcast mechanisms to Xchan:

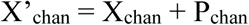

Following coordinate alignment, the matrices are routed into their respective isolated Transformer Encoders (self.temporal_encoder and self.channel_encoder). Each encoder block is configured with a depth of num_layers=2 and utilises num_heads=4 parallel attention heads. The feedforward network multiplier is set to dim_feedforward = hidden_dim × 4 = 512, employing Gaussian Error Linear Unit (GELU) activations and a regularisation dropout of 0.3 (EEGNetEnhanced) to prevent over-parametrisation.

#### 2.9.5 Low-Rank Consensus and Spatiotemporal Broadcast Fusion

The final critical operation of the FactorizedTransformer block is the reconciliation of these decoupled time and space embeddings into a singular, unified diagnostic representation. Because the pathways possess different sequence lengths (T’=128 vs. H=64), they cannot be directly fused via element-wise addition or matrix multiplication.

To address this dimensional mismatch, we fuse the two pathways via low-rank consensus. The temporal feature matrix emerging from the encoder, Ztime ∈ R ^B^ ^×^ ^T’^ ^×^ ^128^, undergoes a global temporal reduction pooling along its sequence axis (dim=1):

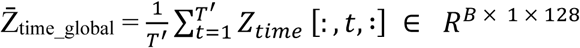

This global average pooling compresses the temporal sequence into a fixed-length summary vector that preserves waveform morphology while discarding phase shifts.

Finally, this global temporal token is broadcast across the spatial sequence axis and added directly to the spatial output matrix Zchan ∈ R ^B^ ^×^ ^H^ ^×^ ^128^ coming out of the channel encoder:

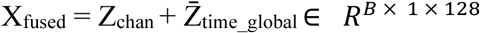

This broadcast fusion scheme establishes a powerful, low-rank additive synthesis. The unified output tensor Xfused overlays the global temporal morphology of the spike onto the localised cross-channel spatial connectivity weights.

#### 2.9.6: Architectural Variants: M2 and M3

To ablate the factorised design, we implemented two single-axis Transformer variants that share the identical front-end (MultiScaleConvBlock) but differ in the back-end aggregation. Variant A replaces the factorised dual encoders with a single Transformer that processes time only (no separate spatial pathway). Variant B makes the same single-axis design deeper (6 layers, 12 heads) to test whether increased model capacity alone can match the factorised design. Full layer-wise configurations, schematics, and dimension tables for these variants are provided in Appendix D. These variants are used solely as ablative controls (Section 3.1) and as complementary components in the ensemble (Section 2.11).

### 2.10 Optimisation Objectives and Ensemble Strategy

#### 2.10.1 Multi-label Optimisation Objective and Hemispheric Anatomical Constraint

Given that the downstream task is formulated as a six-class multi-label classification problem (where each anatomical region is independently predicted as either involved or not involved in the seizure onset network), the choice of the loss function and the incorporation of domain-specific anatomical priors are critical for stable convergence and clinical reliability.

#### 2.10.2 Focal Loss for Handling Sparse Regional Imbalance

As shown in Figure 5, the label distribution is imbalanced, with common temporo-parietal regions and rare occipital regions (∼10%). This poses a training challenge because pure occipital labels are scarce and mostly appear in complex, noisy multifocal epilepsy cases, hindering model adaptation. Furthermore, signals from the deeper parts of the occipital lobe do not always project clearly to standard surface electrodes [40], which exacerbates classification difficulty. Consequently, during early training, the model tends to exploit easier, more frequent regions as a shortcut to achieve a high score.

To mitigate this class imbalance without resorting to computationally expensive resampling, we adopt the Multilabel Focal Loss mechanism [41]. Unlike BCE, Focal Loss introduces a dynamic scaling factor (1−pt)^γ^ to its standard cross-entropy formulation:

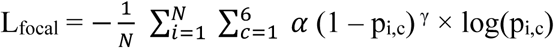

Where N is the batch size, c ∈ {1…,6} indexes the six anatomical regions, p i,c is the model’s predicted probability for sample i and region c, γ=2.0 is the focusing parameter, and αc is the class-specific positive weight (α=3.0 for occipital regions, 1.0 otherwise).

*In our implementation, the focusing parameter is set to γ=2.0. Additionally, to counteract the extreme scarcity of occipital lobe samples, we apply a positive weight factor of α=3.0 to the occipital regions during loss computation, while keeping α=1.0 for all other anatomical regions*

Where p_i,c_ denotes the model’s predicted probability for the i-th sample and the c-th region, and γ(set to 2.0 in our implementation) serves as a focusing parameter. For well-classified, high-confidence samples, (1−pt)^γ^ → 0, effectively down-weighting their contribution to the loss. Conversely, for hard-to-classify or rare samples where pt is small, the scaling factor approaches 1, preserving the full gradient magnitude. These mathematical priors force the network to allocate more representational capacity to the challenging occipital and frontal polar regions, drastically improving the recall rate for low-prevalence focal epilepsies.

#### 2.10.3 Hemisphere Mutual Exclusivity Penalty

While we apply a hard hemispheric constraint at inference time (see Section 2.10.4), we additionally introduce a soft hemispheric penalty during training. This dual approach ensures that (i) the model learns to suppress cross-hemispheric false positives during gradient updates (L_hemi), and (ii) the final output strictly respects unilateral anatomy via post-processing thresholding. The soft penalty during training is critical because it guides the network’s internal representations toward physiologically plausible solutions before the hard constraint is applied at test time.

Although the model processes each window independently as a multi-label task, allowing multiple regions within the same hemisphere to be simultaneously active, consistent with the spatial propagation of interictal discharges, the hemisphere-level penalty enforces unilateral focus at the loss level, aligning with the clinical definition of focal epilepsy. This hybrid formulation preserves within-hemisphere multi-regional discriminability while penalising cross-hemisphere confusion.

To forcibly align the network’s spatial attention with this clinical ground truth, we introduce a novel Hemispheric Mutual Exclusivity Penalty (L_hemi) that is directly injected into the optimisation objective. Let the sigmoid-activated probability vector be denoted as p ∈ R^6^. We define the maximum activated probability for the left and right hemispheres as:

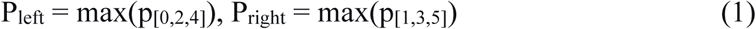

The penalty term is then formulated as the product of these two maxima:

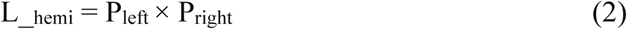

This penalty is minimised (to zero) only when one hemisphere’s maximum probability is zero, aligning with the clinical axiom of unilateral focal epilepsy; it peaks when the model confidently predicts bilateral activity, which is anatomically impossible. Minimising this product during training repulses the probability landscapes of the two hemispheres, guiding the network toward physiologically plausible solutions. The penalty is weighted by λ = 0.1, determined via grid search over {0.05, 0.1, 0.2, 0.5} on the validation set.

#### 2.10.4 Session-Level Binarisation with Anatomical Prior

During inference, after the hemispheric penalty term has guided the network toward unilateral spatial focus during training, we apply a hard anatomical decision rule at the session level. For each continuous recording session, the per-window probability vectors are first averaged to obtain a session-level probability *p* ∈ R^6^. The hemisphere with the higher maximum probability is selected as the putative seizure onset side, and the probabilities of all three regions in the opposite hemisphere are forced to zero:

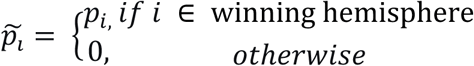

Finally, A fixed binarisation threshold τ = 0.5 (the natural decision boundary of the sigmoid output) is applied to the retained probabilities to produce the final multi-label prediction:

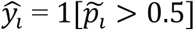

This two-stage decision pipeline, side selection followed by region-level thresholding, ensures that the final output strictly respects the clinical axiom of unilateral focal epilepsy while maintaining fine-grained regional discriminability.

#### 2.10.5 Joint Optimisation Objective

The final combined loss function for end-to-end training integrates the discriminative power of the Focal Loss with the anatomical prior of the Hemispheric Penalty, weighted by a scalar hyperparameter λ:

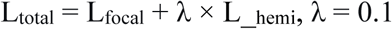

The implementation details of this loss function (including the grid search for λ) are described in Appendix A.

#### 2.10.6 Intra-Architecture Weight Averaging (Model Soup)

To reduce variance from stochastic training dynamics without incurring the inference-time cost of conventional ensembles, we adopt uniform weight averaging (model soup) following Wortsman et al. [42]. For each architecture, we train three independent variants differing only in random seed and mini-batch order. Because these variants converge to proximate regions of the loss landscape, their parameter vectors can be averaged element-wise without performance degradation. Let {θ1,θ2,θ3} denote the checkpoints of the three runs; the fused model is obtained as:

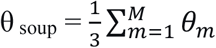

This yields one fused model per seed family. Specifically, we fuse three M1 variants (seeds 100, 101, 102), one M2 variant (seed 42), and one M3 variant (seed 43), totalling five models for the final heterogeneous ensemble. This hierarchical approach eliminates the need to run multiple M1 variants separately at test time while preserving the benefits of multi-run consolidation. Specifically, after the initial training converges, we apply a light fine-tuning stage with three distinct Gaussian noise injection levels (as detailed in the supplementary figure) to the gradients, generating three perturbed checkpoints per converged model. These perturbed weights are then averaged to form the final fused model, further smoothing the loss landscape and improving out-of-distribution robustness.

### 2.11 Heterogeneous Ensemble Strategy

#### 2.11.1 Motivation for Architectural Diversity

While weight averaging consolidates models sharing the same backbone, ensembling structurally distinct models offers complementary inductive biases. Our empirical analysis (Section 3.1) reveals a dissociation: while SOZNet achieves higher global Exact Match, model 1 *shows* superior recall on the rare occipital class at the single-model level. Occipital spikes are characterised by brief, low-amplitude transients that may be better captured by content-based retrieval, whereas temporo-parietal propagation benefits from explicit spatiotemporal factorisation.

Aggregating these structurally distinct models therefore reduces both bias and variance by covering neurophysiological regimes that no single architecture captures equally well. The ensemble thus reduces both bias (via architectural diversity) and variance (via soft voting), yielding a more stable and generalisable decision boundary.

#### 2.11.2 Ensemble Composition

We trained five independent copies of each architecture variant (M1, M2, M3) with different random seeds to select the final ensemble members from the candidate pool. In total, we obtained 15 candidate models. Then we performed an exhaustive search over all non-empty subsets of these 15 candidates on the validation set, with a cardinality cap of five, to maximise exact match rate and recall.

Our final ensemble comprises five constituent models:

- Three M1 variants (seeds 100, 101, 102), each obtained via the model soup procedure described previously. These share the same architecture but diverge due to different random initialisations and fine-tuning trajectories.
- One M2 model (seed 42), fused from its three fine-tuned variants.
- One M3 model (seed 43), fused from its three fine-tuned variants.

This composition balances intra-architecture consolidation (three M1 models capture training stochasticity) with inter-architecture diversity (M2 and M3 introduce distinct structural priors).

#### 2.11.3 Soft Voting via Probability Averaging

At inference time, each constituent model independently processes the same input EEG window X ∈ R^18×2048^ and produces a six-dimensional probability vector p(m) = σ(z(m)) ∈ R^6^, where σ(⋅) denotes the sigmoid function applied elementwise to the raw logits z(m).

The ensemble prediction is obtained via soft voting, i.e., arithmetic averaging of the probability vectors:

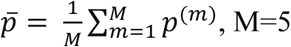

We used soft voting to preserve confidence information. This is particularly critical for scalp EEG, where low-SNR windows produce low-confidence probabilities; soft voting naturally attenuates overconfident but potentially spurious predictions from individual models.

### 2.12 Implementation Details and Evaluation Metrics

#### 2.12.1 Training Configuration

All experiments were implemented in PyTorch and executed on a single NVIDIA RTX 6000 Ada GPU. Models were trained for 30 epochs using the AdamW optimiser (initial learning rate 1 × 10 ^−4^, weight decay 1 × 10 ^−4^) with a warmup-cosine annealing learning rate schedule. Early stopping with a patience of 20 epochs was applied based on validation exact-match accuracy; the checkpoint with the highest validation accuracy was retained for final evaluation. Full hyperparameter details are provided in Appendix A.

A two-stage learning rate schedule was applied to stabilise early training and facilitate fine convergence:

1. Warmup phase (5 epochs): A linear warmup scheduler (LinearLR) increased the learning rate from 1 × 10 ^−6^ (1% of the initial LR) to 1 × 10 ^−4^ over the first 5 epochs.
2. Cosine annealing phase (remaining epochs): Following warmup, a CosineAnnealingLR scheduler decayed the learning rate from 1 × 10 ^−4^ to a minimum of 5 × 10 ^−5^ over the remaining E − 5 epochs, where the total number of epochs E = 30 (config.EPOCHS).

Early stopping with a patience of 20 epochs (config.PATIENCE) was applied based on validation exact-match accuracy; the checkpoint with the highest validation accuracy was retained for final evaluation. The total training time for the largest variant (M3) was approximately 18 hours on the 80-patient training set. Notably, SOZNet achieves this performance with only 0.8M trainable parameters, which is substantially smaller than the 11M-parameter M3, confirming that the factorised design, rather than raw parameter count, drives the performance gain. Baseline models (EEGNet, ShallowConvNet, DeepConvNet, TSCeption) were re-implemented from their original public codebases [18, 19, 43] and trained under the identical data split and hyperparameter tuning protocol as SOZNet (Appendix A). No task-specific modifications were made to these architectures to preserve a fair comparison.

#### 2.12.2 Inference Protocols and Post-Processing

##### We evaluated at two granularities

- Window-level: Each 8-second EEG chunk (18×2048-time steps) was fed independently into the model. The six sigmoid outputs were thresholded at 0.3 (as used in the training code’s preds = (torch.sigmoid(logits) > 0.3).float()) to produce a multi-label prediction for that window. This threshold was chosen empirically to balance precision and recall during early validation. The lower window-level threshold (0.3) compensates for single-window noise. At the session level, averaging across hundreds of windows attenuates transient artefacts, allowing a more conservative threshold of 0.5 without sacrificing sensitivity.
- Session-level: For each continuous recording session, the per-window probability vectors were averaged over all windows belonging to that session. The hemisphere with the higher maximum average probability was selected as the putative seizure onset side; probabilities in the opposite hemisphere were set to zero, and a threshold of 0.5 (config.SESSION_PROB_THRESHOLD) was applied to the remaining three regions to produce the final multi-label prediction. This post-processing step mimics clinical decision-making, where a full session’s worth of data is aggregated.

#### 2.12.3 Evaluation Metrics

We reported the following metrics for both window- and session-level predictions:

- Precision, Recall, and F1-Score: Computed per region (macro-averaged across the six labels) using precision_recall_fscore_support (sklearn) with average=’binary’ per region and zero_division=0.
- Exact Match Ratio (EM Rate): The proportion of samples (windows or sessions) for which all six predicted labels exactly matched the ground-truth labels, calculated via accuracy_score(sklearn).
- Hamming Loss: The fraction of incorrect label predictions, reported for completeness at the session level.

#### 2.12.4 Model Checkpointing and Reproducibility

For each architecture variant, three independent runs were conducted with different random seeds (42, 43, 100, 101, 102). All random seeds were fixed using set_seed() in the training script, covering Python’s random, NumPy, and PyTorch’s CUDA random generators. The best-performing checkpoint (by validation EM rate) was saved for each seed; these checkpoints were subsequently fused via weight averaging (model soup) and then used in the heterogeneous ensemble.

## 3 Results

### 3.0 Overview of Experimental Results

Five key findings emerge from our experiments. First, the ensemble achieved 88.46% session-level Exact Match on the internal test set, outperforming all generic EEG baselines (≤50%) and the best single model (80.77%). Second, systematic ablations confirmed that the factorised spatiotemporal design, rather than model capacity, drove performance, with the full augmentation pipeline contributing a +2.66 p.p. macro-F1 gain and model soup recovering the best single-model performance without inference-time cost. Third, performance generalised robustly across three clinically realistic scenarios: annotation-sparse holdout (82.76% EM), cross-institutional paediatric data (84.62% EM), and mixed with unlabelled random windows (66.67% EM on single model M1). Fourth, mechanistic interpretability was supported by case-level trajectory analysis tracking expert summaries, and by zero false positives on 108 healthy controls. Finally, the entire 24-hour recording pipeline completed in 4.26 minutes, with inference under 8 seconds. Each finding is expanded in the subsections that follow.

### 3.1 Primary Localisation Accuracy: Validating Dataset Adequacy and Baseline Performance

Our primary evaluation focused on the SOZNet-Ensemble, which combines three SOZNet variants, Model 2, and Model 3 through soft voting. On the internal test set (10 patients, 26 sessions). Figure 9 shows that the ensemble achieved an overall Exact Match (EM) ratio of 88.46% (95% CI: 69.8%–97.6%). Across brain regions, the ensemble achieved good recall (1.0) in five out of six regions. The LO region achieved recall of 1.000 but precision of 0.600, resulting in an F1 of 0.750.

**Figure 9:**
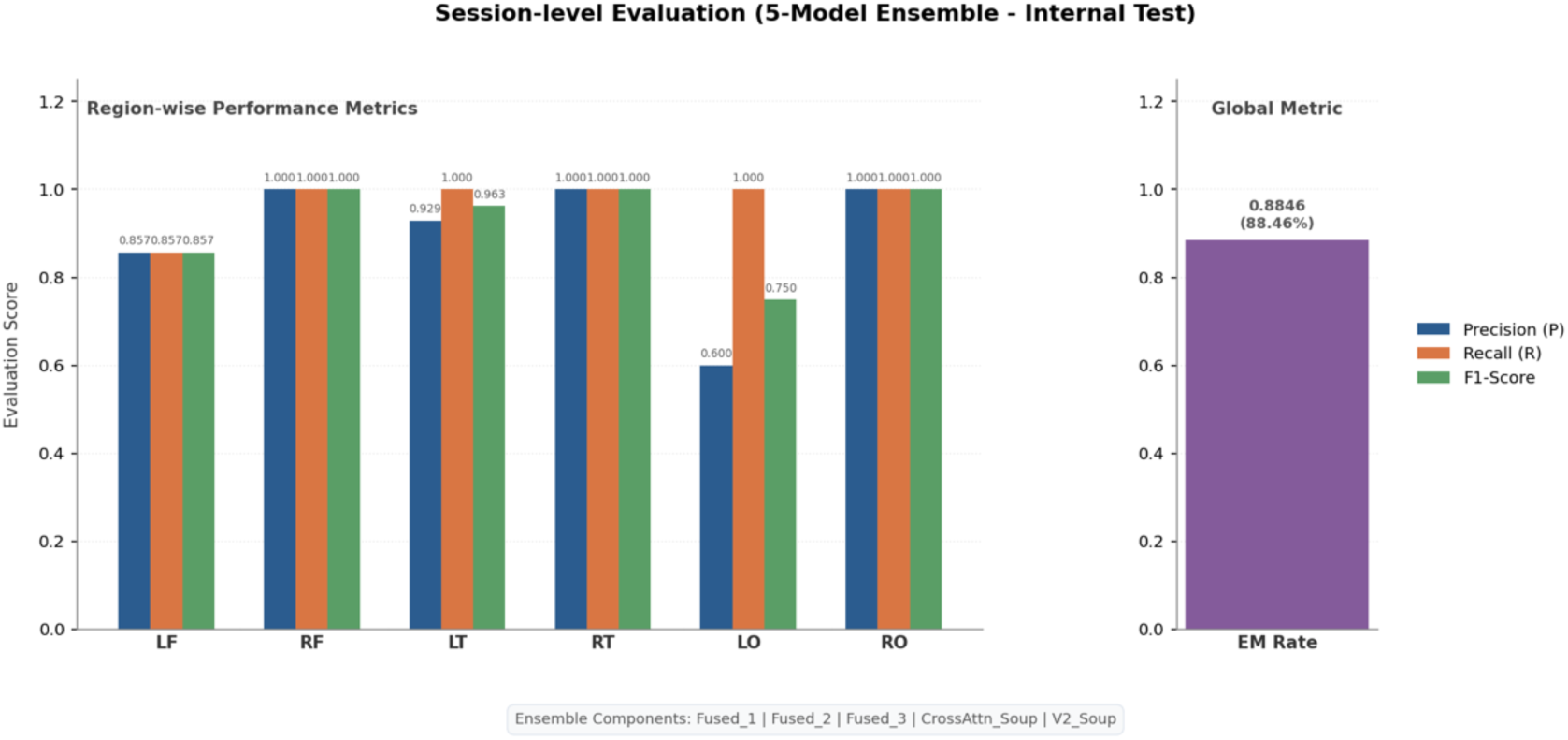
Region-wise precision, recall, and F1 scores of the 5-model ensemble on the internal test set (26 sessions from 10 patients). The left occipital region shows lower precision (0.600) but good recall (1.000); this is a clinically acceptable trade-off, prioritising sensitivity over specificity for epidemiologically rare occipital lobe epilepsy. All other regions achieve F1 ≥ 0.857, with R2, R4, and R6 reaching good scores (1.000); the exact match rate achieved 88.46% with patient reports from epilepsy experts.

For wider comparison, we compared our models against four baseline architectures (Figure 10): EEGNet, ShallowConvNet, TSCeption, and DeepConvNet [18, 43]. As illustrated in Figure 10, all baselines achieved EM ratios below 50.00%. Specifically, our single SOZNet (80.77% EM) outperformed the top-performing baseline, DeepConvNet (50.00%), by an absolute margin of 30.77%, and nearly doubled the performance of TSCeption (46.15%).

**Figure 10:**
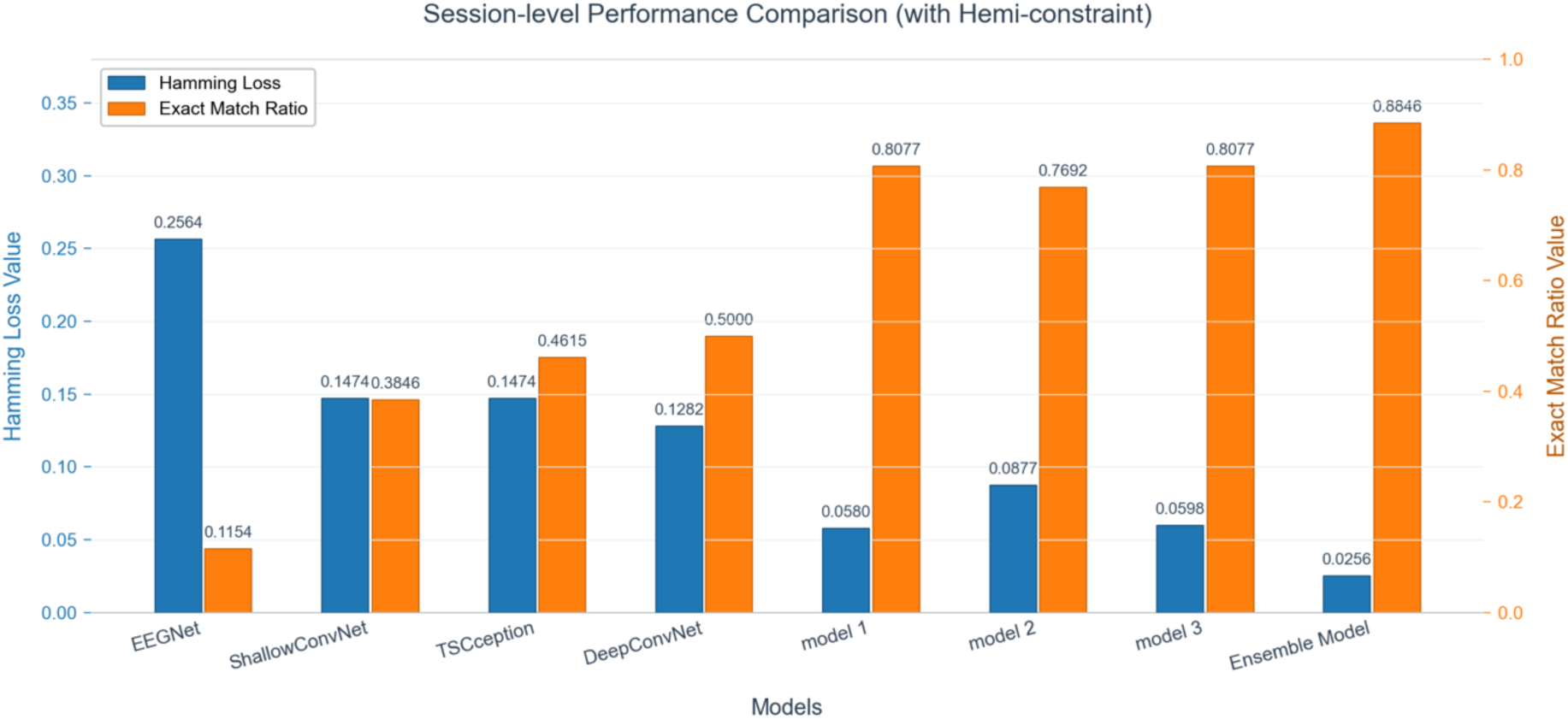
Session-level Exact Match (EM) comparison between the proposed models (single SOZNet and 5-model ensemble) and four generic EEG baselines (EEGNet, ShallowConvNet, TSCeption, DeepConvNet) on the internal test set (26 sessions). Generic baselines fail to exceed 50% EM due to their fixed receptive fields, which cannot simultaneously capture brief spikes (20–70 ms) and longer sharp waves (70–200 ms). The 5-model ensemble achieves 88.46% EM—a 38.31 percentage-point gain over the best baseline (DeepConvNet, 50.00%). Hamming Loss is reduced by an order of magnitude (ensemble: 0.0256 vs. EEGNet: 0.2564). Error bars represent 95% confidence intervals (Clopper-Pearson).

When utilising the heterogeneous 5-model ensemble, the performance gap widened, peaking at 88.46% EM. Additionally, this improvement in full-match accuracy was paired with a significant reduction in error; the ensemble model reduced the Hamming loss to 0.0256, which is an order of magnitude lower than EEGNet’s 0.2564.

We analysed the five constituent models individually (Table 1). All single models achieved high Macro AUCs (≥ 0.960). Integrating these individual variants into the 5-model ensemble improved the overall EM ratio by 7.68%. Although M1-SEED101 achieved a slightly higher Macro AUC (0.989 vs. 0.985), the ensemble produced the highest EM (88.46%, 95% CI: 69.8%–97.6%).

**Table 1:** Session-level performance of individual constituent models and the final 5-model ensemble on the internal test set. EM: Exact Match ratio (percentage of sessions where all six region labels were correctly predicted). CI: 95% confidence interval computed using the Clopper-Pearson exact method. Macro AUC: macro-average of per-region area under the receiver operating characteristic curve. The ensemble achieves the highest EM (88.46%) despite a marginally lower Macro AUC than M1-SEED101 (0.985 vs. 0.989), confirming that ensemble aggregation primarily improves prediction stability and exact-match accuracy rather than ranking performance alone.

| MODEL | EM (%) | 95% CI | MACRO AUC |
| --- | --- | --- | --- |
| **M1(SEED100)** | 76.92 % | [56.4, 91.0] | 0.975 |
| **M1(SEED101)** | 80.77 % | [69.8, 97.6] | 0.989 |
| **M1(SEED102)** | 80.77 % | [60.6, 93.4] | 0.984 |
| **M2(SEED42)** | 76.92 % | [56.4, 91.0] | 0.960 |
| **M3(SEED43)** | 80.77 % | [60.6, 93.4] | 0.980 |
| **ENSEMBLE(5-MODEL)** | 88.46 % | [69.8, 97.6] | 0.985 |

To assess the reliability of the ensemble’s improvement, we performed McNemar’s exact test on paired session-level predictions from the internal test set (n=26) [44]. While the test was underpowered to detect statistical significance at conventional thresholds (two-sided p-values ranged from 0.25 to 0.50) given the small sample size, the asymmetric disagreement pattern was highly consistent: the ensemble never underperformed any single model (n₁₀ = 0 for all comparisons), yet consistently corrected 2–3 misclassified sessions per individual variant (n₀₁ ≥ 2). This consistent direction, combined with a clinically meaningful effect size (7.69 percentage-point gain in EM) and, importantly, the same improvement trend replicated on the larger holdout cohort (n=87, +5.37 p.p.), strongly suggests that the ensemble confers a systematic benefit beyond random variance. We interpret this as evidence of practical superiority, acknowledging that formal statistical confirmation would require a larger prospective test cohort.

### 3.2 Ablation Studies: Validating Architectural Design and Data Augmentation

We ablated the architectural design by comparing M1, M2, and M3. The factorised Transformer (M1, 0.8M parameters) achieved 80.77% EM on the internal test set, outperforming the single-axis Transformer (M2, 3.5M) at 76.92% and matching the deeper single-axis variant (M3, 11M) also at 80.77%. This confirms that architectural factorisation, not raw parameter count, drives performance, consistent with our hypothesis that decoupling temporal and spatial attention mitigates the volume conduction confound. The complementary strengths of M1 (global accuracy) and M2 (occipital sensitivity, recall = 1.000) motivated the heterogeneous ensemble strategy reported in Section 3.1.

We performed systematic ablations on the validation set to quantify each component’s contribution. The full augmentation pipeline (Figure 11) yielded a +2.66 p.p. improvement in macro-F1 over the non-augmented baseline. Gaussian noise (+1.42 p.p.) and hemispheric channel swap (+1.29 p.p.) were the most influential regularisers, which reflects the model’s sensitivity to EMG noise and lateralisation. Random amplitude scaling had the smallest effect (+0.94 p.p.), implying that inter-patient impedance variation has limited impact on performance.

**Figure 11:**
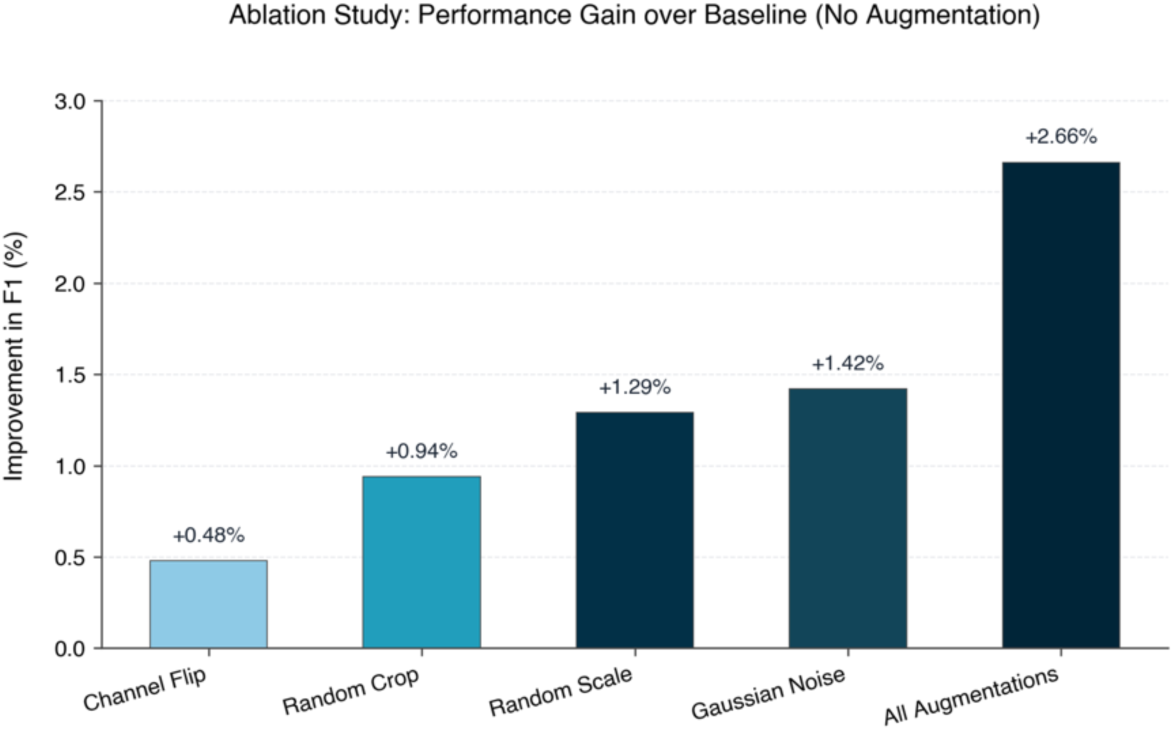
Ablation study of online data augmentation strategies. The bar chart shows the improvement in session-level F1 score on the validation set relative to a baseline model trained without any augmentation. Individual augmentations include Random Crop, Gaussian Noise, Random Amplitude Scaling (“Random Scale”), and Hemisphere Swap, which were evaluated in isolation, alongside the full pipeline (“All Augmentations”). All four strategies contributed positively, with Channel Flip and Random Scale providing the largest individual gains. The full augmentation pipeline gave the highest improvement (+2.66 p.p.), confirming that the combined effect is larger than any single augmentation alone.

Model soup (Figure 12): Uniform weight averaging of three M1 checkpoints raised EM from 76.92% (the best single M1 variant on the validation set) to 80.77%, effectively recovering the performance of the best individual run without incurring any inference-time ensemble cost, with a marginal macro-F1 advantage over greedy soup (0.9094 vs. 0.8927). This indicates that weight averaging smooths the loss landscape and consolidates variance from stochastic training dynamics, providing a stable and computationally efficient alternative for deployment where consistent performance across sessions is more clinically valuable than seed-dependent AUC fluctuations.

**Figure 12.**
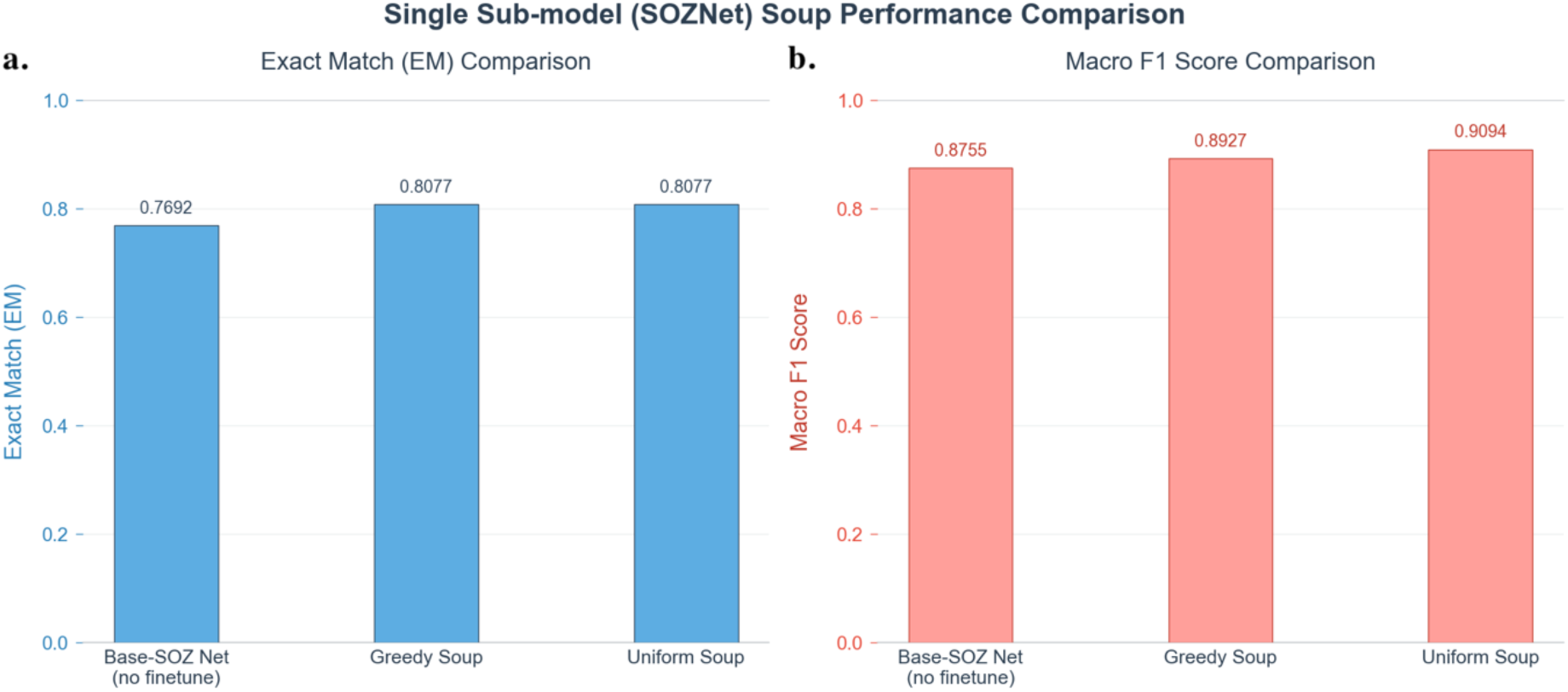
Performance comparison of intra-architecture weight averaging (Model Soup) strategies for the single sub-model (SOZNet). (a) Exact Match (EM) ratios across the baseline SOZNet without fine-tuning (76.92%), Greedy Soup (80.77%), and Uniform Soup (80.77%). (b) Macro F1 scores for the corresponding models, showing an increase from 0.8756 (baseline) to 0.8927 (Greedy Soup) and 0.9094 (Uniform Soup). Both weight averaging strategies demonstrate a numerical improvement in both global match accuracy and region-wise F1 scores compared to the non-fine-tuned baseline.

### 3.3 Generalisation Performance: Validating Clinically Realistic Scenarios

We further evaluated the ensemble on an independent holdout set (87 sessions from 28 patients) and a second independent external test set from Boston Children’s Hospital (BCH), comprising 9 patients with 13 recording sessions from an independent institutional source with different demographics, equipment, and protocols. The ensemble achieved 82.76% EM on the holdout set and 84.62% on the BCH external set (Figure 13), these results suggest cross-institutional generalisability. The 5.70 percentage-point gap between internal (88.46%) and holdout (82.76%) performance was not statistically significant (two-proportion Z-test, Z=0.70, p=0.49), the results from the holdout set support the framework’s generalisability to annotation-sparse scenarios where only interictal spikes are available without corresponding ictal markers.

**Figure 13.**
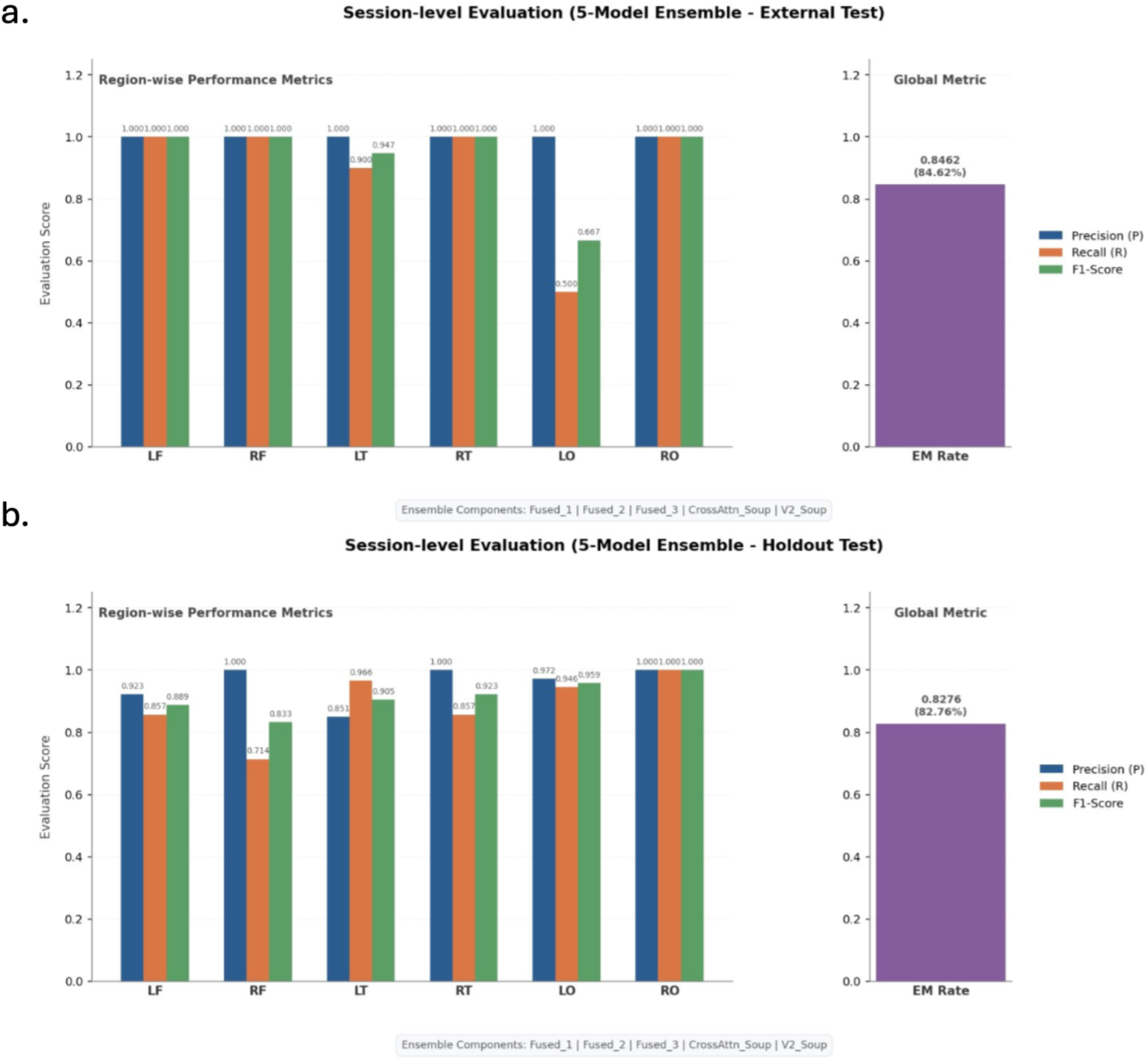
Session-level performance of the 5-model ensemble across the independent holdout and external validation datasets. EM: Exact Match ratio; CI: 95% confidence interval. (a) Model performance on the independent holdout dataset (n = 87 sessions from 28 patients), achieving a global EM rate of 82.76% (95% CI: 73.2%–89.3%). Localised variations are present across brain regions, with the right frontal (RF) region exhibiting a precision of 1.000 and a recall of 0.714. (b) Model performance on the Boston Children’s Hospital (BCH) external test set (n = 13 sessions from 9 patients), yielding a global EM rate of 84.62% (95% CI: 57.8%–95.7%). The BCH cohort entirely comprises paediatric and young adult subjects under 25 years of age, including a subset of 5 patients aged 1 year or younger. A localised drop in sensitivity is observed in the left occipital (LO) region, where recall fell to 0.667.

Performance varied by region across the validation cohorts. In the holdout dataset, the right frontal (RF) region demonstrated precision of 1.000, though its recall decreased to 0.714. Conversely, the left occipital (LO) region, which exhibited lower precision in internal testing, achieved a precision and recall of 1.000 and 0.941, respectively, in the holdout set.

On the BCH external dataset, the model maintained high regional accuracy, particularly in the temporal (LT, RT) and frontal (LF, RF) regions, where both precision and recall reached 1.000. However, a localised reduction in sensitivity was observed in the LO region, where recall fell to 0.667. The age distribution of the BCH cohort spanned entirely under 25 years, including a subset of 5 patients aged 1 year or younger.

In clinical practice, most EEG recordings lack manual spike annotations. To test whether our model can still work in this setting, we ran an additional experiment on the holdout set using completely unlabelled windows, extracted at random intervals without any reference to spike timestamps.

Therefore, an experiment was conducted on the holdout set using unlabelled windows extracted at random intervals without reference to spike timestamps (Figure 14). Operating on these unlabelled windows, the single M1 model achieved an EM ratio of 66.67%, representing a 9.19 percentage-point decrease from the 75.86% achieved on labelled-anchored windows. Both macro-and micro-averaged metrics exhibited slight declines under the unlabelled setting, with the macro-F1 score dropping from 0.870 to 0.831. Region-specific analysis revealed divergent performance trends across different brain areas when manual labels were removed. Specifically, the bilateral frontal regions demonstrated marginal performance improvements in the unlabelled setting. In contrast, the bilateral occipital regions exhibited the highest sensitivity to the absence of annotations, showing substantial decreases in their respective F1 scores. Class 3 remained the largest source of both true positives and false positives under both settings.

**Figure 14.**
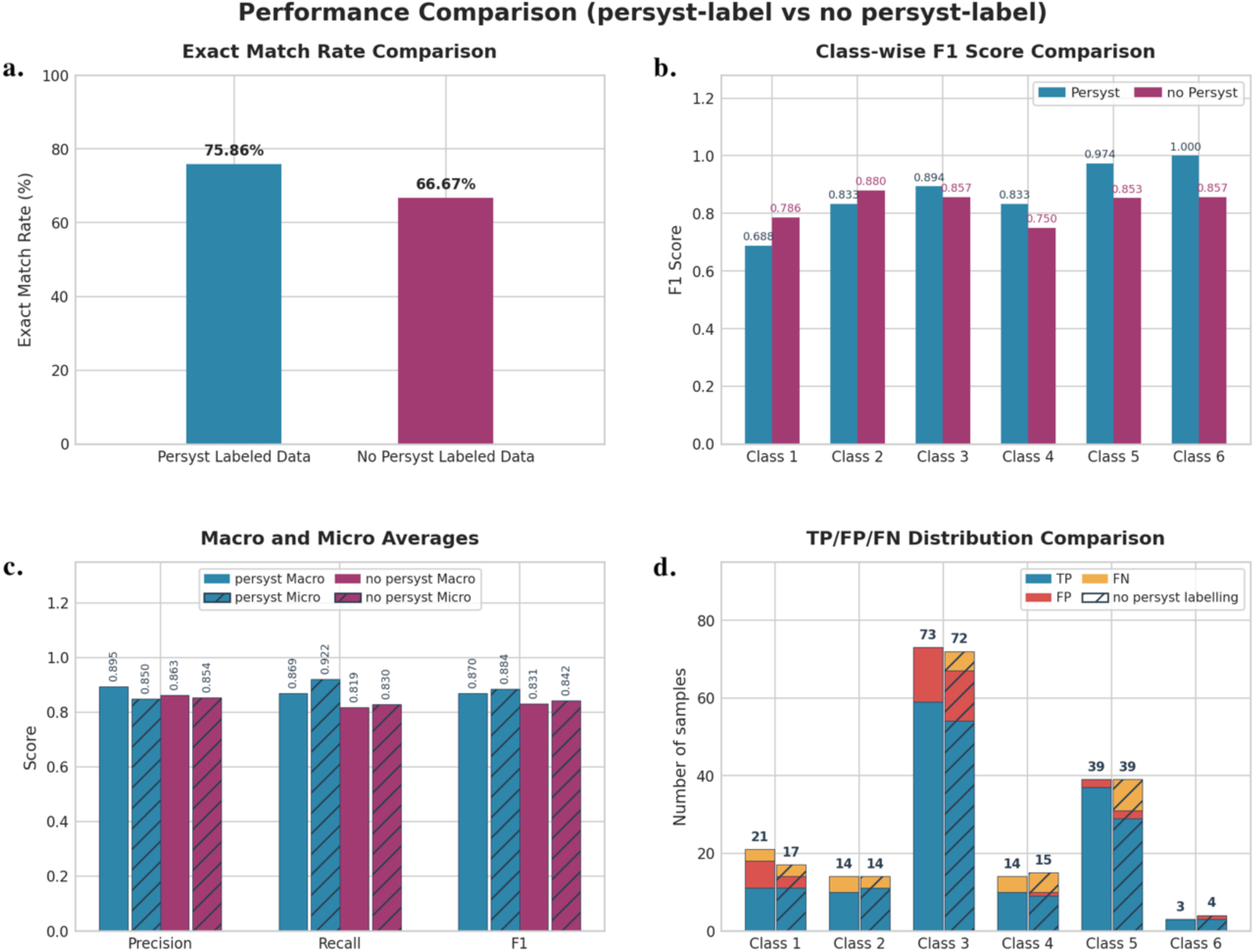
Performance comparison of the single model (M1) on labelled anchored versus randomly extracted unlabelled windows from the holdout set. (a) Exact Match (EM) rates across the two settings, showing a localised decline in the unlabelled condition. (b) Class-wise F1 scores across six anatomical regions, illustrating a divergent trend where bilateral frontal regions (LF, RF) show marginal improvements while bilateral occipital regions (LO, RO) exhibit the most substantial performance drops in the absence of labels. (c) Robustness across macro-and micro-averaged metrics (Precision, Recall, and F1), displaying slight overall decreases under the unlabelled setting. (d) Detailed TP/FP/FN distribution comparison across all brain areas, showing that Class 3 remains the primary source of both true positives and false positives across both validation environments.

We also tested the ensemble on 108 healthy control EEG recordings from PhysioNet [45]. The model produced zero false positives across all subjects, accurately distinguishing common physiological artefacts, including eye blinks, muscle activity, and electrode pops, from true epileptic activity.

### 3.4 Interpretability and Deployment Readiness: Validating Safety and Clinical Utility

#### Case-Level Interpretability

We present two longitudinal case studies to demonstrate the practical clinical utility of our system. By generating continuous probability trajectories over time, the system provides a standardised time index that enables clinicians to trace regional localisation dynamics across long-term scalp EEG recordings and the corresponding confidence. As shown in the profiles of patient 150003760 and patient 150003559 (Figure 15), this allows for the visual verification of the specific continuous data blocks that drive the aggregate SOZ localisation decision.

**Figure 15.**
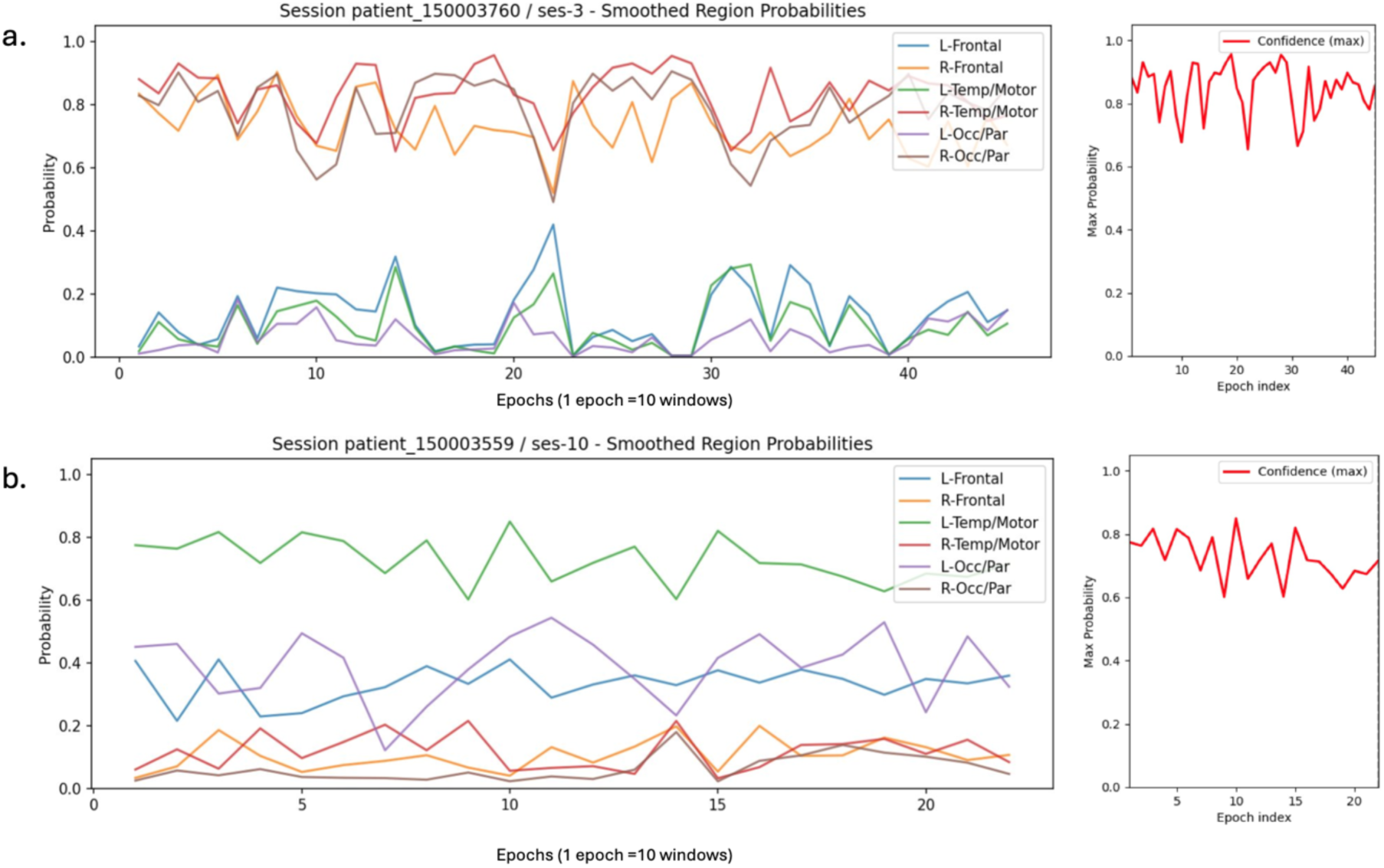
Longitudinal case studies showing smoothed probability trajectories from the test cohort. Probabilities were averaged using a sliding window of 10 consecutive segments to suppress transient single-window noise while preserving underlying network dynamics. The confidence curves (red, right panels) show the maximum prediction probability over time. (a) (Patient 150003760): Consistently high probabilities (>0.80) and confidence (0.65–0.95) are assigned to the right temporal/motor and right frontal regions throughout the recording, which corresponded to the expert summary documenting four electrographic seizures originating from the right temporal region with prominent hemispheric propagation. Around Epoch index 22, right-hemisphere probabilities transiently decrease while the left frontal probability increases, capturing cross-hemispheric propagation noted in the clinical report. (b) (Patient 150003559): Hierarchical distribution with a dominant left temporal/motor trajectory (0.65–0.85, confidence: 0.60–0.85), persistent secondary left occipital/parietal activation (∼0.40), and an intermediate left frontal baseline (∼0.35). This distribution aligns with the complex epileptological profile of a left temporal onset and rare independent bilateral frontal discharges with left predominance (L > R).

The continuous probability trace (Figure 15) assigned consistently high probabilities (>0.80) to both the right temporal/motor and right frontal regions throughout the recording. This output directly mirrored the expert clinical summary, which documented four electrographic seizures originating from the right temporal region, a prominent epileptogenic field spreading across the entire right hemisphere with occasional left-hemisphere involvement, and frequent bursts of disorganised slowing within these exact territories. Notably, during a specific interval around the 22nd recording epoch, the right-hemisphere probabilities experienced a transient downtrend synchronised with a distinct upward spike in the left frontal curve, coinciding with the cross-hemispheric propagation documented in the clinical report.

Case Study 2 (Patient 150003559): Minority Class Isolation. For this patient (Figure 15), the ensemble placed the left temporal/motor region as the primary dominant trajectory (0.65– 0.85), while maintaining a persistent secondary tier for the left occipital/parietal region (∼0.40) and an intermediate baseline for the left frontal region (∼0.35). This hierarchical distribution aligned with the complex epileptological profile: one electroclinical seizure with left temporal onset, frequent high-voltage epileptiform discharges in the left temporal region, and rare independent discharges in the frontal regions bilaterally with higher density on the left (L > R). The model identified the primary left temporal focus, isolated the adjacent occipital/parietal boundary, and distinguished the subtle bilateral frontal asymmetry.

Finally, we benchmarked the end-to-end processing pipeline on a full 24-hour continuous unlabelled EDF recording across three sampling rates. The total execution time was 4.26 minutes at 256 Hz, 7.26 minutes at 512 Hz, and 10.76 minutes at 1024 Hz (Figure 16). Execution profiling revealed that data ingestion and signal harmonisation (including EDF parsing, filtering, and resampling) consumed over 82% of the total runtime. In contrast, deep learning model inference required less than 8 seconds under all tested configurations, indicating that neural network computation is not the primary performance bottleneck of the pipeline. For context, a 24-hour EEG recording typically requires 60–120 minutes of expert visual review to identify and annotate epileptiform events [18]. The current pipeline’s 4.26-minute processing time represents a 14-to 28-fold acceleration over manual review, even in the unlabelled scenario where no spike timestamps are available. This comparison is not intended to suggest replacement of clinical expertise but to quantify the potential for workflow triage in high-volume epilepsy centres: a tool that can process an overnight recording before the morning ward round could flag high-probability cases for early expert review.

**Figure 16:**
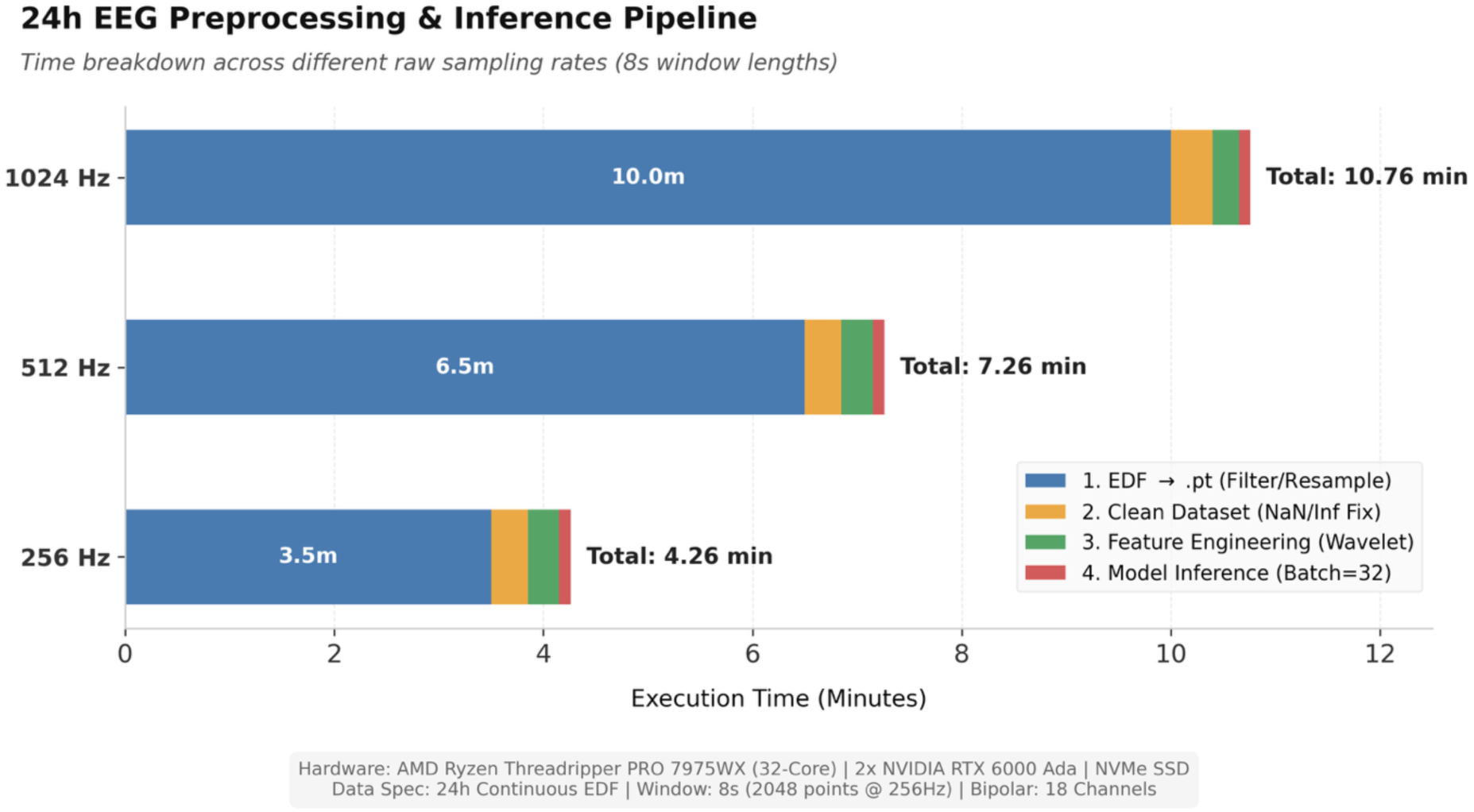
End-to-end processing time for a full 24-hour continuous EEG recording. Measurements were performed on a system with AMD Ryzen Threadripper PRO 7975WX (32-core), 2× NVIDIA RTX 6000 Ada GPUs, and NVMe SSD storage. Left panel: Total processing time across three sampling rates: 4.26 minutes at 256 Hz, 7.26 minutes at 512 Hz, and 10.76 minutes at 1024 Hz (8-second window length, 18-channel bipolar montage). Right panel: Runtime breakdown by pipeline stage at 256 Hz. Data ingestion and signal harmonisation (EDF parsing, filtering, and resampling) consume >82% of total runtime, while deep learning model inference completes in under 8 seconds across all configurations. This profile confirms that the neural network is no longer the computational bottleneck, making the framework suitable for high-throughput clinical screening operations.

## 4: Discussion

### Interpretation of Results

Our results suggest a broader point: the spatial information needed for SOZ localisation survives the spatial blurring of volume conduction, provided the model respects EEG’s physical structure. That is, temporal dynamics and spatial propagation should be processed through separate pathways, with anatomical priors explicitly encoded in the loss function. Three observations support this interpretation. First, the factorised architecture (M1, 0.8M parameters) matched the performance of a much deeper single-axis variant (M3, 11M), indicating that architectural inductive bias matters more than raw capacity. Second, the hemispheric penalty reduced anatomically implausible bilateral predictions without harming within-hemisphere discriminability. Third, the model’s zero false-positive rate on healthy controls suggests it learned propagation dynamics rather than simple waveform templates -a point we return to below. With this framing, we turn to the detailed findings.

The ensemble model achieved 88.46% session-level exact match (EM) on the internal test set, much higher than all baseline models (≤50%). Single lightweight Model 1 maintained 66.67% EM on random windows with a non-spike labelling setting. The ensemble model also produced zero false positives on 108 healthy controls, and processed a full 24-hour recording in 4.26 minutes. These results indicate that routine scalp EEG can provide lobar SOZ localisation, extending prior evidence from invasive [46] and high-density [47] recordings to the low-density, low-SNR clinical setting.

Standard baseline architectures rely primarily on localised convolutional layers with fixed receptive fields. Deep variants such as DeepConvNet stack layers to capture abstract representations, but successive downsampling sacrifices fine temporal resolution and fails to preserve transient paroxysmal features such as brief spikes (20–70 ms). EEGNet is shallow but lacks the capacity to bridge long-range temporal dependencies. Our architecture combines parallel multi-scale convolutions with subsequent Transformer layers. The multi-scale convolutions preserve both frequency components and sharp transients, while the self-attention mechanism enables non-local temporal modelling without loss of structural resolution.

Within this framework, the smaller Model 1 (0.7M) performed similar with the deeper Models 2 (3M) and Model 3 (11M), providing insights into spatiotemporal regularisation. Models 2 and 3 use standard Transformer encoders on an unflattened spatiotemporal grid, incurring quadratic complexity (O(N²)) and potential overfitting on sample-constrained clinical data. Model 1 instead uses a factorised architecture that separates attention into distinct spatial and temporal dimensions. Physically, this factorisation reflects the orthogonal properties of scalp EEG: temporal dynamics reflect neural synchrony, while spatial dynamics reflect volume conduction and topographic distribution. By encoding this inductive bias, the factorised transformer reduces parameter redundancy and improves generalisation.

Our results indicate that these architectures have complementary feature extraction abilities. The factorised transformer (Model 1) is tailored for canonical spatiotemporal landmarks, while the deeper standard transformers (Models 2 and 3) better capture unconstrained global network dynamics. Their prediction errors are largely uncorrelated. Soft voting averages their continuous probability distributions, reducing idiosyncratic false positives or false negatives. This aggregation yielded a 7.69 percentage-point gain in EM (from 80.77% to 88.46%) over the best single variant.

Comparing the ensemble against individual sub-models using McNemar’s exact test on paired session-level predictions (Section 3.1) yielded two-sided p-values of 0.25 and 0.50, both above the 0.05 threshold. This reflects the limited sample size (n = 26 sessions). McNemar’s test depends on the frequency of discordant pairs; with few discordant sessions, statistical power is low regardless of the percentage improvement. The ensemble consistently matched or outperformed all individual models, with zero negative discordant pairs (n₁₀ = 0). It corrected single-model errors in 2–3 sessions, raising EM by 7.69 percentage points. Random variance would produce a symmetric distribution (n₁₀ ≈ n₀₁), so this asymmetry supports systematic error correction. This pattern is consistent with the generalisation metrics on the larger holdout and multi-centre cohorts.

The performance contributions of different augmentations (Section 3.2) provide insights into model regularisation. Gaussian noise injection yielded the largest individual improvement (Macro F1 +1.42%). This complements the preprocessing denoising procedures. Preprocessing filters (e.g., bandpass and notch filtering) remove macro-level, high-amplitude artefacts; training-time stochastic noise injection simulates micro-level residual variations, such as subtle fluctuations in electrode-skin impedance and hardware thermal drift that standard filters cannot fully eliminate. Random Gaussian perturbation encourages the model to learn signal rather than noise, improving robustness in unlabelled test settings.

The optimisation objective incorporated two domain-specific priors. Focal Loss (γ = 2.0) with class-specific positive weighting (α = 3.0 for occipital regions) was introduced to counteract the epidemiological under-representation of occipital lobe epilepsy (8.2–8.7% of positive instances). This weighting directs the gradient toward rare classes and helps the ensemble achieve recall of 1.000 for the right occipital region. The hemispheric mutual exclusivity penalty (λ = 0.1), by contrast, enforced a different prior: unilateral focal epilepsy. Its primary effect was not to improve overall accuracy but to suppress anatomically implausible bilateral predictions. Together, these two components operated at different levels: focal loss addressed inter-regional class imbalance, while the hemispheric penalty enforced inter-hemispheric anatomical plausibility.

In the expanded holdout dataset, the global EM rate remained stable (82.76%), but right frontal (RF) recall decreased to 0.714. This drop likely reflects a shift in data distribution: the internal test set (26 sessions) primarily captured dense ictal periods, whereas the holdout cohort (87 sessions) introduced longer continuous recordings with lower discharge density. The inclusion of annotation-sparse windows may increase variance and reduce per-session discharge density, leading to localised sensitivity loss. Nevertheless, RF precision remained 1.000, indicating that positive predictive boundaries stayed reliable.

Performance heterogeneity was more evident in the unlabelled test on randomly extracted windows (Figure 14). Macro-averaged F1 decreased from 0.870 to 0.831, with varied regional effects. Marginal improvements occurred in bilateral frontal regions (LF, RF), while the largest drops were observed in bilateral occipital regions (LO, RO). These differences likely reflect cortical physiology. Frontal lobe discharges propagate broadly and appear frequently in random windows, allowing the multi-scale convolutions to learn regional features. In contrast, occipital paroxysms are typically transient, focal, and state-dependent (e.g., eye-closure or visual suppression) [48]. These brief events are easily masked by background alpha rhythms and physiological noise and thus depend more on structured temporal anchors.

The model achieved 84.62% EM on the external dataset, suggesting cross-institutional generalisability. This validation cohort had a specific demographic profile: all subjects were under 25 years, including five infants aged ≤1 year. Regional accuracies remained high, particularly in temporal and frontal regions, indicating that the learned representations are robust to age-dependent morphological differences in infant cranial structure and developing EEG backgrounds. However, left occipital (LO) recall was lower at 0.667. Although this suggests regional susceptibility, the small BCH sample size (n = 13 sessions from 9 patients) is a clear statistical limitation. The localised LO sensitivity drop may also reflect physical differences in infant EEG recording: open fontanelles and higher CSF-to-brain volume ratios alter volume conduction properties [49], potentially attenuating occipital signals more than frontal or temporal sources. Larger multi-centre trials are needed to validate these regional variations.

The longitudinal case studies (Figure 15) provide a framework for evaluating temporal interpretability. For Patient 150003760, the model produced a continuously elevated probability trajectory (>0.80) for the right temporal and right frontal regions, which corresponded to the clinical report documenting four electrographic seizures originating from the right temporal structures. At the 22nd smoothed epoch index, the right-hemisphere probability curve transiently decreased while the left frontal trajectory showed a reciprocal increase. This crossover coincided with a sharp reduction in prediction confidence (Figure 15, upper right panel). When the discharge remained localised to the primary right temporal focus, the model output a highly certain regional classification. However, during propagation across the corpus callosum to the contralateral hemisphere, the multi-scale convolutions registered conflicting epileptiform features from both sides. The resulting spatial ambiguity flattened the softmax distribution, causing the confidence metric to dip. This alignment between reduced confidence and documented cross-hemispheric propagation supports the model’s ability to produce time-indexed network transitions without relying on post-hoc saliency maps.

The zero false-positive rate on 108 healthy control recordings is noteworthy. Generic transient detectors, whether rule-based or learned, often flag high-amplitude artefacts such as eye blinks, muscle activity, and electrode pops. Our model did not, despite lacking explicit artefact rejection modules. This suggests that the factorised spatiotemporal attention mechanism distinguishes true epileptiform spikes from benign events, likely by learning propagation dynamics. The distinction makes clinical sense: benign artefacts, however large in amplitude, lack the organised spatial propagation patterns that characterise true cortical discharges. Eye blinks project diffusely but without the evolving field topography of an epileptic spike; muscle activity is spatially circumscribed and transient. By learning this topological signature rather than a waveform template, the model approximates a clinical rule of thumb that spikes are defined by field distribution and propagation trajectory [50].] That a purely data-driven model captures this contextual distinction without explicit feature engineering suggests that the ensemble learns network-level dynamics rather than isolated waveform templates, extending the intracranial evidence of Jiang et al. [11] to the noisier regime of routine scalp EEG.

Clinical deployment also requires computational efficiency. Evaluation on a 24-hour EDF recording (Figure 16) showed that end-to-end execution scales linearly with data volume and sampling rate. Runtime profiling shows that data ingestion, signal harmonisation, and hardware operations (EDF parsing, filtering, downsampling) consume over 82% of execution time. In contrast, the 5-model ensemble inference phase requires under 8 seconds for the full 24-hour sequence. Thus, neural network computation is not the primary bottleneck; throughput is limited by CPU-bound data I/O and signal processing. Total processing time remains under 11 minutes at the highest density (1024 Hz), indicating that the framework operates faster than real-time and could serve as an automated screening tool for long-term video-EEG data.

### Limitations

Label granularity and causal limitations. A key limitation is the spatial resolution of the ground-truth labels. Our six-region mapping, which combines temporal and parietal lobes, was a pragmatic alignment with the 18-channel bipolar montage. This coarse labelling presents a ceiling for supervised learning: the model may learn electrophysiological features that contain sublobar information, but it is never incentivised to express them during training. Similarly, the model learns statistical associations between EEG patterns and lobar labels derived from clinical reports, not direct measurements of the epileptogenic zone. It may pick up on correlated features that are not themselves causative, such as secondary propagation patterns or co-occurring structural abnormalities visible on MRI. Prospective comparison with surgical outcomes, the definitive gold standard, would be required to establish causal validity.

Data-related limitations. Occipital lobe epilepsy is an inherent minority in our cohort, reflecting its epidemiological rarity rather than a sampling bias. While Focal Loss and ensemble aggregation improved sensitivity for this rare class, this optimisation reflects a small absolute number of test-set samples. Consequently, these findings should be treated as hypothesis-generating regarding rare-event learning rather than definitive proof. Dedicated occipital-specific cohorts would be required for prospective validation.

Clinical translation limitations. The framework is positioned as a decision-support tool for use after routine scalp EEG acquisition, prior to sEEG implantation planning. Its output is a six-region probability vector that requires interpretation by an experienced epileptologist; it is not a standalone diagnostic. The current implementation is retrospective; prospective validation in real-time clinical settings is essential.

The inference-time hemispheric hard constraint assumes confirmed focal epilepsy and is not designed to differentiate focal from generalised cases. However, we tested the model on a small subset of sessions with documented bilateral or multifocal onsets—cases explicitly excluded from training. In these instances, the ensemble rarely produced a false-positive unilateral localisation. Instead, probabilities across all six regions remained below the 0.5 threshold, yielding a “no prediction” output. When inspecting the longitudinal probability trajectories, regions consistent with the bilateral SOZ showed elevated probabilities (typically 0.3–0.45) relative to other regions, while completely healthy controls (PhysioNet) produced uniformly low probabilities across all regions (typically <0.15). This suggests that the model’s probability magnitude encodes graded information about the presence of epileptiform activity, even when the unilateral assumption is violated. Clinicians could therefore interpret uniformly low probabilities across all regions as a flag for inconclusive or non-focal cases, rather than accepting a forced lateralisation as valid.

We excluded ambiguous cases, bilateral onsets, and genetic epilepsies from this proof-of-principle study; generalising to these more complex presentations will require further work.

### Future Directions

First, the label granularity limitation identified above could be addressed through teacher-student distillation. A teacher model trained on high-density EEG (128 channels) or ESI source imaging could generate soft spatial probability maps over individual anatomical parcels. These maps could then serve as supervisory signals for distilling sublobar discriminative features into our low-density 18-channel architecture, potentially recovering finer-grained information that our current six-region labels cannot express.

Second, the dissociation between M1 and M2 performance on occipital versus temporal classes suggests that no single architectural prior is optimal for all epilepsy subtypes. A subtype-adaptive ensemble could address this: rather than a fixed heterogeneous ensemble, one could train specialised models for different electroclinical syndromes (e.g., temporal lobe epilepsy vs. occipital lobe epilepsy) and route inputs to the most appropriate model based on preliminary spectral features. This builds directly on our observation that M1 captures sustained spatiotemporal propagation patterns while M2 better detects sparse, transient discharges. Such an approach would require larger, subtype-stratified cohorts but could improve performance across the full spectrum of focal epilepsies.

Third, the factorised attention mechanism provides a basis for interpretable source localisation. Unlike standard transformers where attention weights are difficult to interpret due to flattened time-space sequences, our architecture produces separate temporal and spatial attention maps. Projecting spatial attention weights onto the 10–20 electrode topology could generate a dynamic topographical heatmap showing, in real time, which electrode clusters the model is attending to, providing clinicians with visualisation directly mappable to the EEG montage.

Fourth, the computational profile-less than 8 seconds of inference versus over 4 minutes of data I/O for a 24-hour recording-suggests that model compression is not the primary engineering priority. Instead, developing streaming EDF parsers and edge-AI deployment could enable continuous, window-wise inference directly on ICU or EMU bedside devices. Coupled with an automated summariser, this could transform the pipeline from a batch screening tool into a live decision-support system that flags high-confidence localisations without waiting for full-recording download.

Finally, integrating multimodal imaging could generate more precise spatial ground truths. ESI has demonstrated the ability to localise averaged interictal discharges to within 10 mm of the surgical resection margin in up to 67% of patients with favourable outcomes [51]. Instead of training on coarse lobar labels, one could project EEG source localisation results onto individual structural MRI scans and use the resulting 3D coordinates as continuous or fine-grained anatomical targets, leveraging EEG’s temporal precision and MRI’s spatial resolution.

In conclusion, this work demonstrates proof-of-principle that a deep learning framework can reliably localise the seizure onset zone at the lobar level from routine 18-channel scalp EEG. We curated and annotated a large, clinically heterogeneous cohort of 100 patients (262 sessions, >6,000 hours), and introduced a targeted online augmentation pipeline—Gaussian noise, amplitude scaling, temporal cropping, and hemispheric channel swap—that mitigated overfitting with systematic ablations confirming each component’s contribution. The factorised spatiotemporal Transformer, combined with a hemispheric mutual exclusivity penalty and a heterogeneous five-model ensemble, achieved 88.46% session-level exact match on the internal test set, outperforming generic EEG baselines by over 38 percentage points. Performance generalised to annotation-sparse holdout data (82.76%), a cross-institutional paediatric cohort (84.62%), and wholly unlabelled random segments (66.67% with single M1), with case-level trajectory analysis confirming that predictions track expert clinical summaries across continuous recordings. Critically, by offering a non-invasive, computationally efficient (4.26 minutes for a 24-hour recording), and interpretable pathway to expedite presurgical evaluation, this approach has the potential to reduce reliance on prolonged inpatient monitoring and invasive implantation, thereby lowering patient risk and accelerating surgical decision-making for those with drug-resistant focal epilepsy.

## Declaration of Generative AI Assistance

DeepSeek-R1 was used as a functional tool during the preparation of this thesis. It was utilised solely to assist with preliminary structural planning and the grammatical refinement and polishing of my original text. The AI did not generate any academic content, arguments, or data analyses, and the final work remains entirely my own.

## Appendix A: Training Hyperparameters

### Content A1: Complete Training Configuration Table

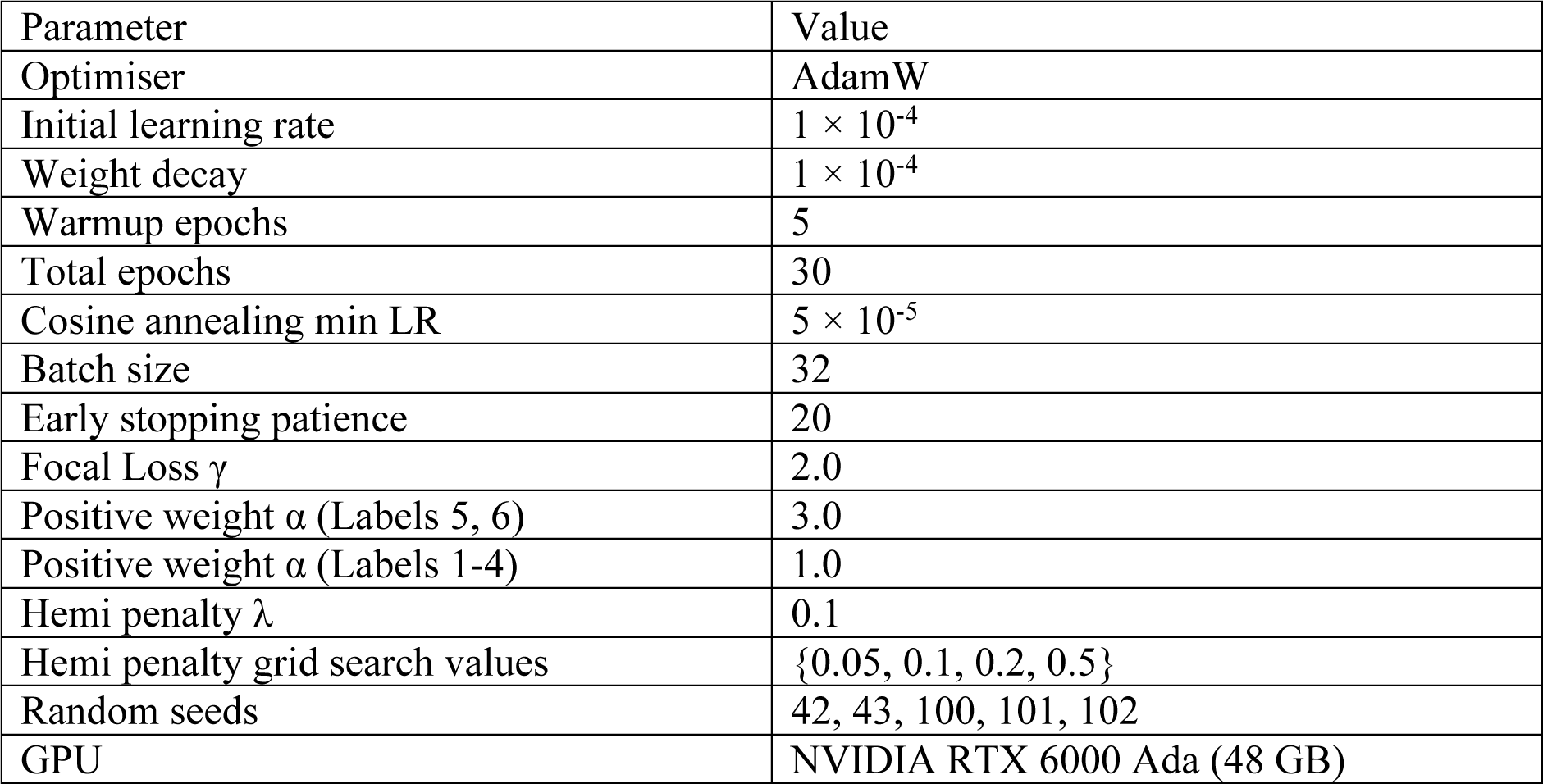

| Parameter | Value |
| --- | --- |
| Optimiser | AdamW |
| Initial learning rate | $1 \times 10^{-4}$ |
| Weight decay | $1 \times 10^{-4}$ |
| Warmup epochs | 5 |
| Total epochs | 30 |
| Cosine annealing min LR | $5 \times 10^{-5}$ |
| Batch size | 32 |
| Early stopping patience | 20 |
| Focal Loss $\gamma$ | 2.0 |
| Positive weight $\alpha$ (Labels 5, 6) | 3.0 |
| Positive weight $\alpha$ (Labels 1-4) | 1.0 |
| Hemi penalty $\lambda$ | 0.1 |
| Hemi penalty grid search values | {0.05, 0.1, 0.2, 0.5} |
| Random seeds | 42, 43, 100, 101, 102 |
| GPU | NVIDIA RTX 6000 Ada (48 GB) |

### A2: Explanation of the α weight in the loss function

Focal Loss Positive Weighting. The α values were selected based on the inverse class frequency in the training set. Labels 5 (Left Occipital) and 6 (Right Occipital) represented 8.2% and 8.7% of positive instances, respectively; an α of 3.0 was applied to these rare classes to amplify their gradient contribution. All other regions (Labels 1-4) used α=1.0.

### A3: λ Grid Search Results

Hemispheric Penalty Coefficient Search. The coefficient λ was tuned on the validation set over {0.05,0.1,0.2,0.5}. λ=0.1 yielded the highest validation EM (78.85%), compared with76.92% (λ=0.05), 76.92% (λ=0.2), and 73.08% (λ=0.5). All subsequent experiments use λ=0.1.

## Appendix B: Data Preprocessing and Quality Control Details

### B1: Parallel Processing Architecture

To manage the computational overhead of processing long-term continuous recordings, a multiprocessing pipeline was implemented using Python’s multiprocessing module with a dedicated CPU worker pool. Epochs were batched into chunks of 100 segments and distributed across worker processes. Each worker independently performed resampling, trimming, and serialisation. Memory management was enforced via explicit del statements and gc.collect() at the end of each session loop to prevent memory leaks.

### B2: Quality Control Thresholds

-Maximum consecutive NaN: 512 time steps (2 seconds at 256 Hz)
-Global NaN ratio: 30% of the total 18×4096 matrix (>22,118 NaN values)
-Interpolation method: 1D linear interpolation per channel
-Boundary handling: Nearest-neighbour extrapolation for NaN values at sequence start/end

### B3: Dimension Standardisation

The original EEG recordings varied in sampling rate (256 Hz, 512 Hz, and 1024 Hz in the BIDMC cohort). All windows were resampled to 256 Hz using scipy.signal.resample, yielding 4,096 time steps for a 16-second window. The final tensor shape was 1×18×4096 (batch dimension × channels × time steps).

## Appendix C: Signal Processing Parameters

### C1: Wavelet Denoising Configuration

-Mother wavelet: Daubechies 4 (db4)
-Decomposition level: 4
-Threshold method: Universal threshold (VisuShrink) [38]
-Noise standard deviation estimation: Median absolute deviation (MAD) of the finest detail coefficients (d1):

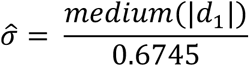

Universal threshold:

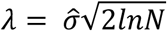

where N is the signal length.

Soft-thresholding function:

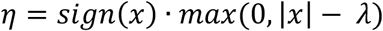

Reconstruction: Inverse discrete wavelet transform (IDWT) after thresholding.

### C2: Common Average Reference (CAR)

After bipolar montage derivation, a spatial CAR filter was applied:

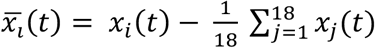

Where x_i_(t) is the voltage at channel i at time t, and the mean is computed across all 18 channels. This removes residual common-mode noise shared across all electrodes.

## Appendix D: Architecture Details of Ablation Variants

To ablate the factorised design in SOZNet (Section 2.9), we introduced two single-axis Transformer variants that share the identical front-end (MultiScaleConvBlock and depthwise spatiotemporal convolution) but differ in the back-end Transformer aggregation. Their full layer-wise configurations are provided below; a schematic comparison is shown in Figure S1.

### D.1 SOZNet (Factorised Spatiotemporal Transformer)

The core architecture is described in Section 2.9. Table D1 provides the layer-wise dimensions.

**Table D1:** SOZNet — Layer-wise Dimensions.

| Layer | Input Shape | Output Shape |
| --- | --- | --- |
| Input | B×1×18×2048 | -- |
| MultiScaleConvBlock<br>(4branches) | B×1×18×2048 | B×32×18×2048 |
| Concatenation | -- | B×32×18×2048 |
| Depthwise Spatial Conv | B×32×18×2048 | B×64×18×2048 |
| Temporal Downsampling<br>(×16) | B×64×18×2048 | B×64×18×128 |
| Squeeze | B×64×18×128 | B×64×128 |
| Temporal Projection | B×64×128 | B×128×128 |
| Spatial Projection | B×64×128 | B×64×128 |
| Temporal Encoder (2 layers,<br>4 heads) | B×128×128 | B×128×128 |
| Spatial Encoder (2 layers, 4<br>heads) | B×64×128 | B×64×128 |
| Global Temporal Pooling | B×128×128 | B×1×128 |
| Broadcast Fusion | B×64×128 + B×1×128 | B×64×128 |
| Classification Head | B×64×128 | B×6 |

Total trainable parameters: 0.8M

### D.2 Model 2 (Single-Axis Baseline)

Instead of factorised dual encoders, Model 2 adopts a single-axis Transformer operating on the temporal sequence with a learnable query for content-based pooling. Table D2 provides the layer-wise dimensions.

**Table D2:** Model 2 — Layer-wise Dimensions.

| Layer | Input Shape | Output Shape |
| --- | --- | --- |
| Input (from MultiScaleConvBlock) | $B \times 64 \times 128$ | |
| Permute | $B \times 64 \times 128$ | $B \times 128 \times 64$ |
| Linear Projection | $B \times 128 \times 64$ | $B \times 128 \times 256$ |
| Trainable Positional Encoding | $B \times 128 \times 256$ | $B \times 128 \times 256$ |
| Transformer Encoder (4 layers, 8 heads) | $B \times 128 \times 256$ | $B \times 128 \times 256$ |
| Learnable Query Cross-Attention | -- | $B \times 1 \times 256$ |
| Classification Head | $B \times 256$ | $B \times 6$ |

Total trainable parameters: 3.5M

### D.3 Model 3 (Deep-Only Baseline)

Model 3 scales the single-axis design to 6 layers with 12 heads, using fixed sinusoidal positional encoding. Table D3 provides the layer-wise dimensions.

**Table D3:** Model 3 — Layer-wise Dimensions.

| Layer | Input Shape | Output Shape |
| --- | --- | --- |
| Input (from MultiScaleConvBlock) | $B \times 64 \times 128$ | -- |
| Permute | $B \times 64 \times 128$ | $B \times 128 \times 64$ |
| Linear Projection | $B \times 128 \times 64$ | $B \times 128 \times 384$ |
| Sinusoidal Positional Encoding | $B \times 128 \times 384$ | $B \times 128 \times 384$ |
| Transformer Encoder (6 layers, 12 heads) | $B \times 128 \times 384$ | $B \times 128 \times 384$ |
| Learnable Query Cross-Attention | -- | $B \times 1 \times 384$ |
| Classification Head | $B \times 384$ | $B \times 6$ |

Total trainable parameters: 11M

### D.4 Schematic Comparison

**Figure S1:**
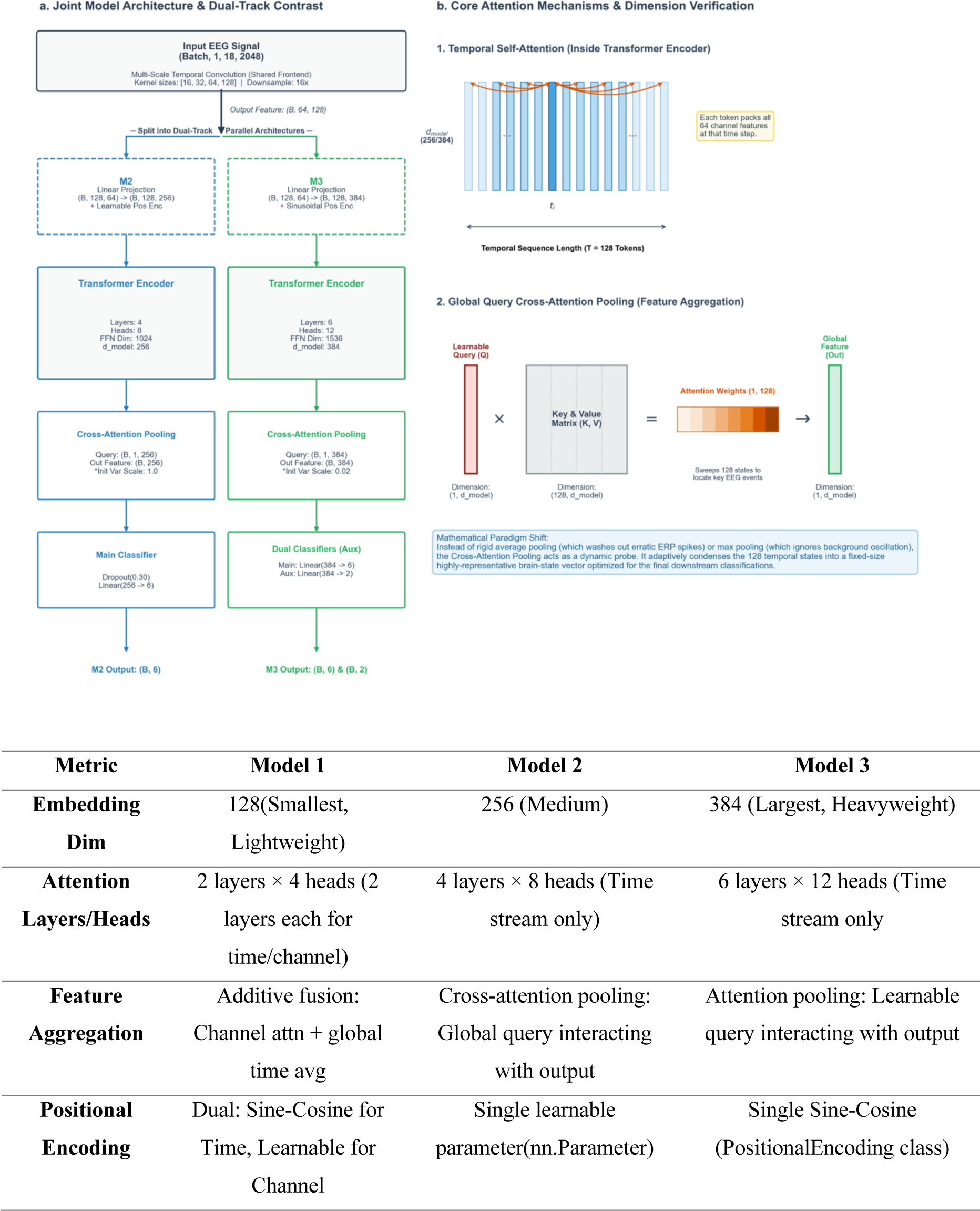
Architectural comparison of Model 2 and Model 3. Both variants share the identical multi-scale convolutional front-end (MultiScaleConvBlock + depthwise spatiotemporal convolution) and classification head, differing only in the back-end Transformer aggregation strategy. Model 2 (top) employs a single-axis Transformer with trainable positional encoding and a learnable global query vector (q∈R ^256^) for content-based cross-attention pooling. Model 3 (bottom) scales to a deeper configuration (6 layers, 12 heads, d model = 384) with fixed sinusoidal positional encoding and more aggressive dropout (0.35). Both use attention pooling with a learnable query.

